# Epha2, an epithelial cell pattern recognition receptor for fungal β-glucans

**DOI:** 10.1101/135434

**Authors:** Marc Swidergall, Norma V. Solis, Scott G. Filler

## Abstract

Oral epithelial cells discriminate between pathogenic and non-pathogenic stimuli, and only induce an inflammatory response when they are exposed to high levels of a potentially harmful microorganism. The pattern recognition receptors (PRRs) in epithelial cells that mediate this differential response are poorly understood. Here, we demonstrate that the ephrin type-A receptor 2 (EphA2) is an oral epithelial cell PRR that binds to exposed β-glucans on the surface of the fungal pathogen *Candida albicans*. Binding of *C. albicans* to EphA2 on oral epithelial cells activates signal transducer and activator of transcription 3 (Stat3) and mitogen-activated protein kinase signaling in an inoculum-dependent manner, and is required for induction of a pro-inflammatory and antifungal response. Inhibition of EphA2 in mice decreases IL-17 signaling during oropharyngeal candidiasis, resulting in increased oral fungal burden and fungal dissemination. Our study reveals that EphA2 functions as PRR for β-glucans that senses epithelial cell fungal burden and is required for the maximal mucosal inflammatory response to *C. albicans.*

**One Sentence Summary:** EphA2 is a pattern recognition receptor that senses fungal β-glucans to induce an inflammatory response in oral epithelial cells.

---

Oral epithelial cells are continuously exposed to a multitude of antigens derived from food and the resident bacterial and fungal microbiota. They must be able to discriminate between pathogenic and non-pathogenic stimuli so that an inflammatory response is induced only when the epithelial cells are exposed to high levels of a potentially harmful microorganism. The interactions of oral epithelial cells with the fungus *Candida albicans* provides an example of this paradigm. As part of the normal microbiota of the gastrointestinal and reproductive tracts of healthy individuals (*1*), this organism elicits a minimal epithelial cell response during commensal growth, when a low number of organisms is present. However, when local or systemic host defenses are weakened, *C. albicans* can proliferate and cause oropharyngeal candidiasis (OPC), an infection that is highly prevalent in patients with HIV/AIDS, diabetes, and iatrogenic or autoimmune-induced dry mouth (*2*). When OPC occurs, oral epithelial cells are activated to induce a pro-inflammatory response that plays a central role in limiting the extent of infection. For example, mice with an oral epithelial cell-specific defect in IL-17 receptor signaling or production of β defensin 3 are highly susceptible to OPC and unable to resolve the infection (*3*).

How oral epithelial cells discriminate between *C. albicans* when it grows as a commensal organism versus an invasive pathogen is incompletely understood. The fungus can interconvert between ovoid yeast and filamentous hyphae. *C. albicans* yeast are poorly invasive and weakly stimulate epithelial cells to release proinflammatory cytokines and host defense peptides (HDPs). In contrast, hyphae avidly invade epithelial cells and strongly stimulate the production of cytokines and HDPs (*4-6*). Mucosal epithelial cells express a variety of pattern recognition receptors (PRRs) that can potentially recognize *C. albicans* (*7*). These cells also express non-classical receptors such as the epidermal growth factor receptor (EGFR), HER2, and E-cadherin that can recognize *C. albicans* hyphae (*8-10*). However, relatively little is known about PRRs in oral epithelial cells, even though these cells constitute a key barrier to mucosal infection.

The ephrin type A receptor 2 (EphA2) is a receptor tyrosine kinase that induces both endocytosis and cytokine production by host cells (*11, 12*). We investigated the hypothesis that EphA2 functions as an epithelial cell receptor for *C. albicans*. By confocal microscopic imaging of intact epithelial cells infected with *C. albicans*, we observed that EphA2 accumulated around yeast-phase cells after 15 min of infection and hyphae after 90 min (Fig. 1A). To ascertain whether *C. albicans* activated EphA2, oral epithelial cells were infected with yeast-phase *C. albicans* cells and the extent of EphA2 autophosphorylation was analyzed over time. EphA2 phosphorylation increased above basal levels within 15 min post-infection, when the organisms were still in the yeast phase (Fig. 1B; S1). EphA2 phosphorylation also remained elevated after 60 and 90 min of infection, when the organisms had formed hyphae. When epithelial cells were incubated for 15 min with either yeast-or hyphal-phase organisms, Epha2 phosphorylation was stimulated to the same extent, indicating that both forms of the organism can activate the receptor (Fig. 1C, S1).

**Fig. 1.**
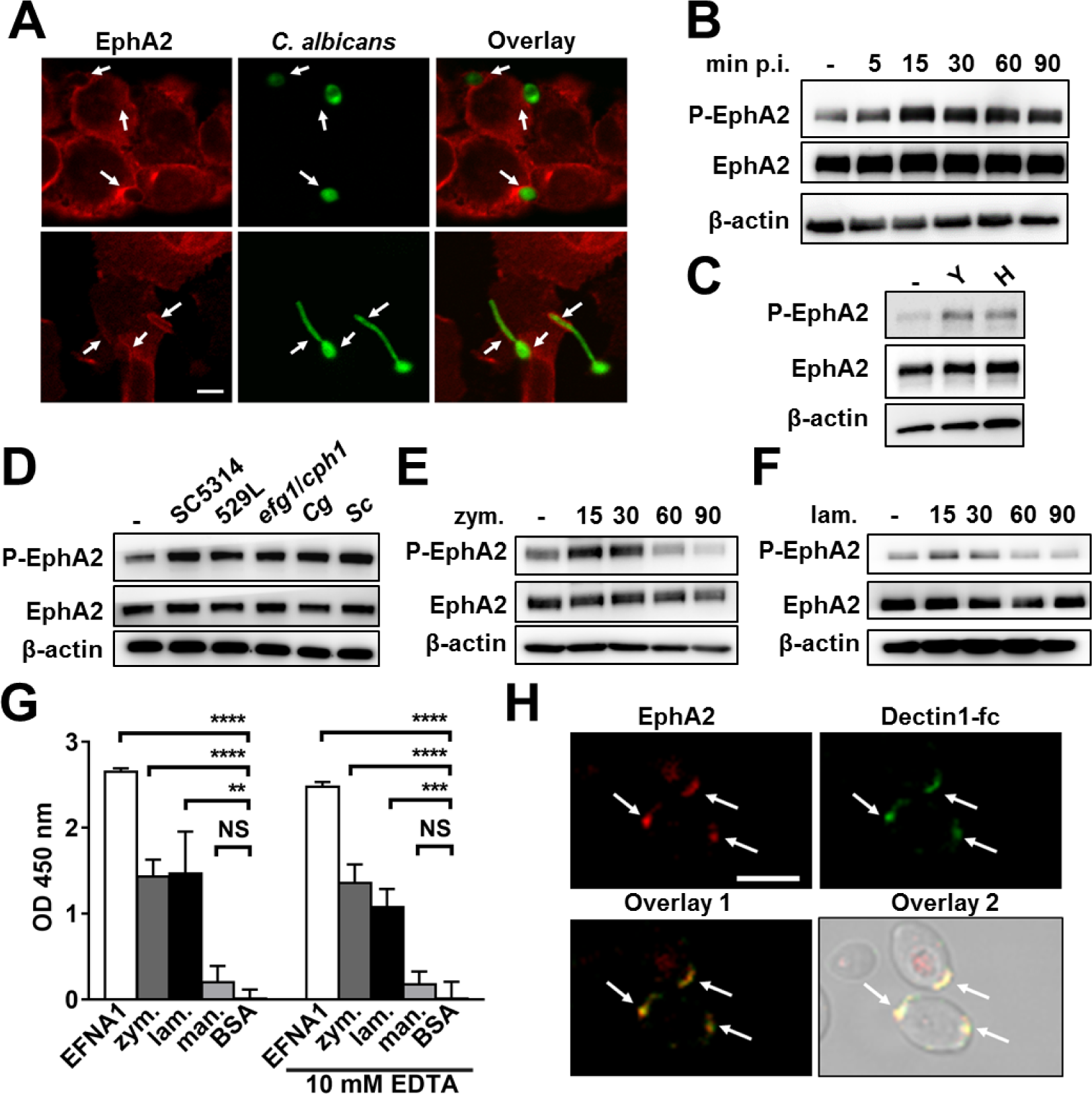
EphA2 is bound and activated by β-glucans. (**A**) Confocal microscopic images of OKF6/TERT-2 epithelial cells that had been infected with GFP expressing *C. albicans* (CAI4-GFP) and then stained for EphA2 (red). Arrows indicate the accumulation of EphA2 around fungal organism. (**B**) Immunoblot analysis showing the time course EphA2 phosphorylation in oral epithelial cells infected with yeast-phase *C. albicans* SC5314 for the indicated times. (**C**) EphA2 phosphorylation after 15-min infection with either *C. albicans* yeast or pregerminated hyphae. H, hyphae; Y, yeast. (**D**) Effects of *C. albicans* (SC5314, 529L, *efg1/cph1*), *Candida glabrata*, and *Saccharomyces cerevisiae* on EphA2 phosphorylation. Cg, *C. glabrata*; Sc, *S. cerevisiae*. (**E-F**) Time course (in minutes) of EphA2 phosphorylation induced by zymosan (**E**) and laminarin (**F**). (**G**) Binding of recombinant EphA2 to immobilized ephrin A1, zymosan, laminarin, mannan, and BSA as determined by ELISA. Results are mean ± SD of 3 experiments, each performed in triplicates. Statistical analysis of binding is shown relative to wells coated with BSA. EFNA1, ephrin A1; lam, laminarin; man, mannan; zym, zymosan (**H**) EphA2 (red) and dectin-1-Fc (green) bind to exposed β-glucan on yeast-phase *C. albicans* organism. \**P* < 0.05, \*\**P* < 0.01, \*\*\**P* < 0.001, \*\*\*\**P* < 0.0001; NS, not significant. Scale bars 5 μm.

To determine the specificity of EphA2 signaling, we analyzed whether bacteria (*Staphylococcus aureus*, *Escherichia coli*), a clinical mucosal *C. albicans* isolate (529L), a yeast-locked *C. albicans* mutant (*efg1*/*cph1*), and different fungal species (*Candida glabrata*, *Saccharomyces cerevisiae*) were able to induce EphA2 phosphorylation. Although neither *S. aureus* nor *E. coli* stimulated the phosphorylation of EphA2 (Fig. S2), all fungi tested induced phosphorylation of this receptor within 15 min of infection (Fig. 1D, Fig. S3), suggesting that EphA2 is activated by a conserved fungal cell wall component. To test this possibility, we analyzed EphA2 activation by β-glucan and mannan, which are present in the cell walls of many fungi. Both particulate β-glucan (zymosan) and soluble β-glucan (laminarin) stimulated EphA2 phosphorylation after 15 and 30 min, and this phosphorylation returned to basal levels by 60 min (Fig. 1E, F, S4). Mannan, another cell wall component, failed to induce detectable EphA2 phosphorylation (Fig. S5).

To ascertain whether EphA2 interacts directly with β-glucans, we used an ELISA to analyze the binding of recombinant EphA2 to potential ligands. As expected, EphA2 bound with high affinity to its natural ligand, ephrin A1 (EFNA1) (Fig. 1G). EphA2 also bound avidly to zymosan and laminarin, but not to mannan in both the presence and absence of 10 mM EDTA, indicating that binding to β-glucans is independent of Ca^2+^. Consistent with the ELISA data, we found that recombinant EphA2 bound to intact yeast-phase *C. albicans* cells at regions of exposed β-glucan, as demonstrated by co-localization with dectin-1-Fc (Fig. 1H). EphA2 also bound to exposed β-glucan on other fungal pathogens, including *Aspergillus fumigatus* and *Rhizopus delemar* (Fig. S6).

*C. albicans* hyphae can invade oral epithelial cells by receptor-induced endocytosis (*10*), and epithelial cell invasion is associated with fungal-induced epithelial cell damage (*6*). We analyzed the effects of the small molecule tyrosine kinase inhibitor, dasatinib (DAS) (*13*), and a specific EphA2 antagonist, 4-(2,5-dimethyl-1H-pyrrol-1-yl)-2-hydroxybenzoic acid (ANT) (*14*) on *C. albicans* invasion and damage of epithelial cells to determine the biological significance of EphA2 activation. After verifying that DAS and ANT inhibited *C. albicans*-induced phosphorylation of EphA2 (Fig. S7), we found that both inhibitors significantly reduced endocytosis of *C. albicans* (Fig. 2A) and host cell damage (Fig. 2B), but had no effect on *C. albicans* growth rate or hyphal elongation (Table S1). Depletion of epithelial cell EphA2 by siRNA similarly inhibited fungal endocytosis and host cell damage (Fig. S8), while addition of EFNA1 to further stimulate EphA2 enhanced both invasion and damage (Fig. 2A, B). Thus, the extent EphA2 activation governs epithelial cell endocytosis of *C. albicans* and subsequent fungal-induced damage.

**Fig. 2.**
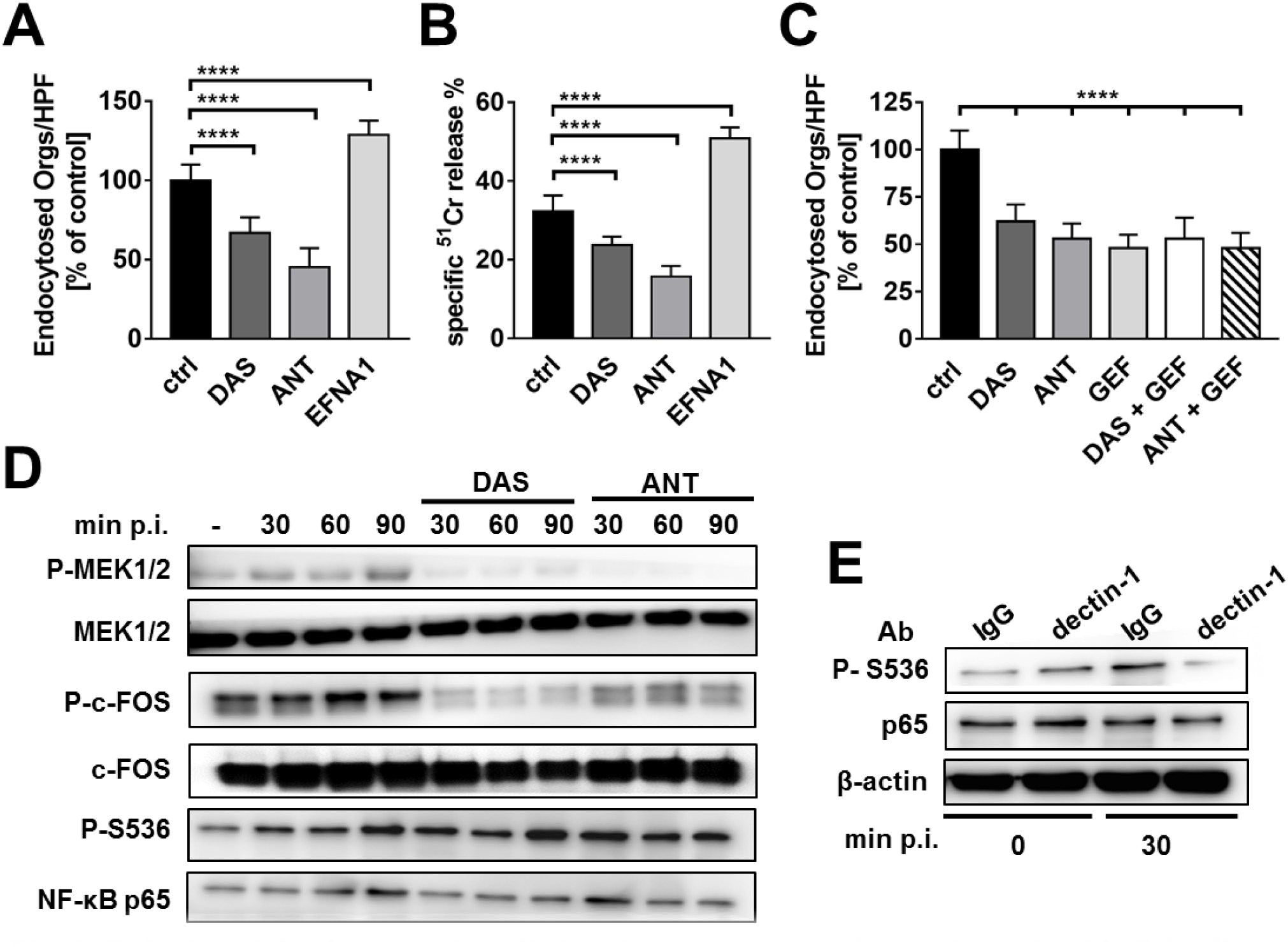
EphA2 and dectin-1 regulate distinct host response pathways in oral epithelial cells. (**A-B**) Effects of treatment of oral epithelial cells with dasatinib, an EphA2 antagonist, or EFNA1 on the endocytosis of *C. albicans* (**A**) and extent of *C. albicans*-induced host cell damage (**B**). Results are the mean ± SD of three experiments, each performed in triplicate. (**C**) Effects of inhibition of EphA2 and/or the epidermal growth factor (EGFR) on the endocytosis of *C. albicans* by oral epithelial cells. Results are the mean ± SD of three experiments, each performed in triplicate. (**D**) Immunoblots showing the phosphorylation of epithelial cells MEK1/2, c-FOS, and S536 of NF-κB p65 in response to *C. albicans* infection in the presence and absence of DAS or ANT. (**E**) Effect of treatment of epithelial cells with an anti-dectin-1 mAb on the phosphorylation of p565 S536 induced by *C. albicans.* ANT, EphA2 antagonist; DAS, dasatinib; GEF, gefitinib; p.i., post-infection. \*\*\*\**P* < 0.0001.

Epidermal growth factor receptor (EGFR) signaling is crucial for inducing the endocytosis of *C. albicans* by oral epithelial cells (*8, 15*). We analyzed the relationship between EGFR and EphA2 signaling during the endocytosis of *C. albicans* and found that inhibition of EphA2 with either DAS or ANT, and inhibition of EGFR with gefitinib (*8*) reduced the endocytosis of *C. albicans* by a similar amount (Fig. 2C). Simultaneous inhibition of both EphA2 and EGFR did not decrease endocytosis further. Also, inhibition of EphA2 with DAS or ANT blocked *C. albicans*-induced EGFR phosphorylation (Fig. S9). Collectively, these results suggest that EphA2 and EGFR function in the same pathway and that EphA2 activation is required for induction of EGFR signaling and subsequent endocytosis of *C. albicans.*

Contact with *C. albicans* yeast cells activates multiple signaling pathways in oral epithelial cells including mitogen-activated kinase regulated kinase (MEK) 1/2, p38, and NF-κB (*16*). We investigated whether EphA2 might function as a receptor that activates these signaling pathways. We determined that *C. albicans* stimulated the phosphorylation of MEK1/2 and p38-mediated phosphorylation of the c-Fos transcription factor within 30 min of infection (Fig. 2D and S10). The phosphorylation of these proteins was maintained for at least 90 min and was inhibited by both DAS and ANT. Although *C. albicans* stimulated phosphorylation of the p65 subunit of NF-κB, this phosphorylation was not inhibited by DAS or ANT, suggesting that NF-κB activation is induced by a receptor other than EphA2. One candidate receptor is the dectin-1 β-glucan receptor, which we determined was expressed on the surface of both the OKF6/TERT-2 oral epithelial cell line and desquamated human buccal epithelial cells (Fig. S11). We found that a neutralizing dectin-1 mAb inhibited *C. albicans*-induced activation of p65 (Fig. 2 E, S12), but had no effect on the phosphorylation of EphA2 (S13). These results suggest that in response to *C. albicans* infection, dectin-1 is necessary for induction of NF-κB signaling in oral epithelial cells, whereas EphA2 is required for activation of the MEK1/2 and p38 signaling pathways.

To investigate EphA2 signaling induced by β-glucan alone, we analyzed the response of epithelial cells to zymosan. While zymosan strongly activated MEK1/2 phosphorylation at 15 and 30 min, phosphorylation returned to basal levels by 60 min (Fig. S14). Also, zymosan did not induce detectable c-Fos phosphorylation over the 90-min duration of the experiment (Fig. S12). Thus, fungal factors in addition to β-glucan are required to sustain MEK1/2 signaling and induce c-Fos phosphorylation.

By secreting proinflammatory cytokines and host defense peptides (HDPs), oral epithelial cells are vital for limiting fungal proliferation during OPC (*17, 18*). The production of many of these factors is governed by the transcription factor signal transducer and activator of transcription 3 (STAT3) (*17, 19*). We hypothesized that the interaction of *C. albicans* with EphA2 in oral epithelial cells would activate Stat3 and stimulate production of proinflammatory cytokines and HDPs. To test this, we determined the effects of DAS and ANT on Stat3 phosphorylation in epithelial cells. By both immunoblotting and ELISA, we found that *C. albicans* stimulated Stat3 phosphorylation within 15 min of infection and that blocking EphA2 with DAS or ANT reduced *C. albicans*-induced Stat3 phosphorylation to basal levels (Fig. 3A, B, S15). Depletion of EphA2 in oral epithelial cells with siRNA also decreased *C. albicans*-induced Stat3 phosphorylation (Fig. S15). Although exposure to zymosan induced Stat3 phosphorylation within 15 min, this phosphorylation was transient, returning to basal levels after 30 min (Fig. 3C, Fig. S15). Inhibition of dectin-1 did not reduce zymosan-induced activation of Stat3 (Fig. S16), suggesting that EphA2 is the primary β-glucan receptor that activates this transcription factor.

**Fig. 3.**
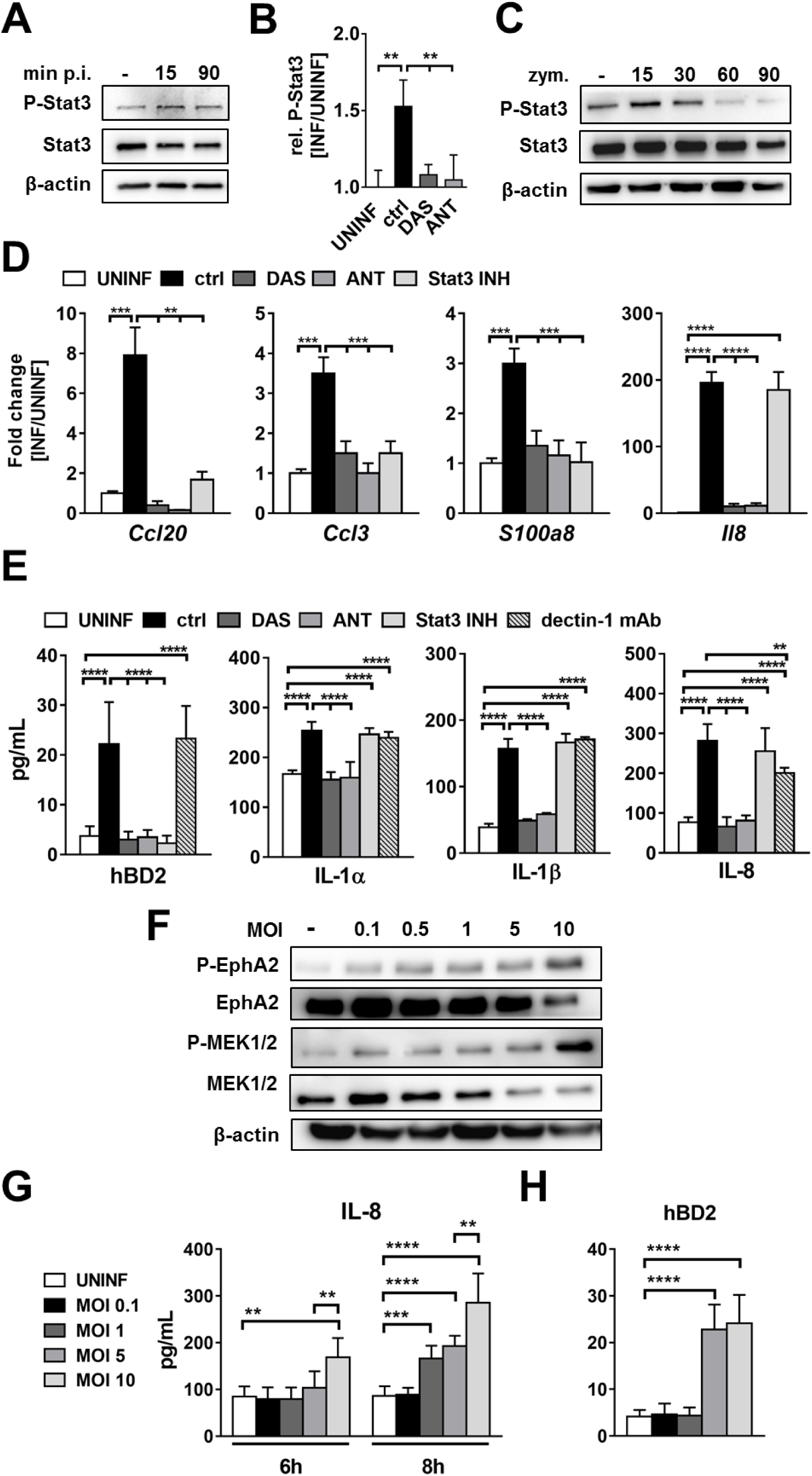
EphA2 signaling regulates the inflammatory response. (**A**) Immunoblot demonstrating that *C. albicans* infection induces phosphorylation of Stat3. (**B**) Effects of the EphA2 inhibitors, DAS and ANT on *C. albicans*-induced phosphorylation of Stat3, as measured by ELISA. Results are mean ± SD of 3 experiments, each performed in 3. (**C**) Time course of Stat3 phosphorylation induced by zymosan. (**D**) Inhibition of EphA2 signaling decreases the epithelial cell pro-inflammatory response to *C. albicans* infection. Oral epithelial cells were incubated with inhibitors of EphA2 and Stat3 and then infected with *C. albicans* for 5 h after which chemokines and the S100a8 alarmin mRNA levels were determined by real-time PCR. Results are mean ± SD of 2 experiments, each performed in triplicates and are presented as fold induction relative to uninfected epithelial cells. (**E**) Inhibition of EphA2 signaling reduces secretion of hBD2, IL-1 and IL-8 induced by 8 h of infection. Results are mean ± SD of at least three independent experiments, each performed in duplicate. Statistical analysis is shown relative to epithelial cells infected with *C. albicans* in the absence of inhibitors. (**F**) Immunoblots showing the effects of increasing *C. albicans* inoculum on the extent of EphA2 phosphorylation. (**G-H**) Effects of *C. albicans* inoculum on epithelial cell secretion of IL-8 (**G**) and hBD2 (**H**). Results are the mean ± SD of 3 independent experiments in duplicates. ctrl, control; hBD2, human β defensin 2; MOI, multiplicity of infection; Stat3 INH, Stat3 inhibitor; UNINF, uninfected. \**P* < 0.05, \*\**P* < 0.01, \*\*\**P* < 0.001, \*\*\*\**P* < 0.0001; NS, not significant.

By blocking EphA2 signaling with DAS, ANT or the Stat3 inhibitor S31-201 (*20*), we found that the EphA2-Stat3 axis regulates *C. albicans*-induced mRNA expression of the chemokines *Ccl20* and *Ccl3*, and the alarmin *S100a8 in vitro* (Fig. 3D). It also governs the secretion of human β-defensin 2 (hBD2) (Fig. 3E and S17). Although EphA2 inhibition reduced *Il8* mRNA expression and secretion of IL-8, IL-1α and IL-1β, inhibition of Stat3 did not (Fig. 3D, E). Thus, these responses are induced by EphA independently of Stat3. Treatment of epithelial cells with DAS or ANT did not inhibit production of IL-8 induced by TNF-α and IL-17 (Fig. S18), indicating that these inhibitors had no effect on the epithelial cell response to a stimulus other than β-glucan.

The addition of zymosan to the epithelial cells did not stimulate secretion of IL-8 or hBD2 (Fig. S19), suggesting that although β-glucan-induced EphA2 activation is required to prime epithelial cells, a second signal is necessary to prolong EphA2 activation and stimulate cytokine and HDP release. Dectin-1, another β-glucan receptor (*21, 22*), played limited role in the epithelial cell response to *C. albicans*; an anti-dectin-1 antibody had no effect on the release of hBD2, IL-1α, or IL-1β by infected epithelial cells, although it did cause a modest reduction in IL-8 release (Fig. 3E).

We investigated whether the extent of EphA2 activation governed the epithelial cell response to infection with different inocula of *C. albicans*. We found that while there was a gradual increase in the extent of EphA2 phosphorylation as the multiplicity of infection (MOI) increased from 0.1 to 5 organisms per epithelial cell, EphA2 phosphorylation dramatically increased at a MOI of 10 (Fig. 3F, S18). Low level phosphorylation of MEK1/2 was also stimulated by *C. albicans* yeast at a MOI of 0.1 to 1, whereas phosphorylation increased exponentially at MOIs of 5 and 10 (Fig. 3F, S18).

Phosphorylation of Epha2 was blocked by DAS and ANT at all MOIs (Fig. S18). The increased phosphorylation of EphA2 and MEK1/2 at high MOIs corresponded to enhanced secretion of IL-8 and hBD2 (Fig. 3H, I). Thus, while low inocula of *C. albicans* induce modest EphA2 activation and a minimal inflammatory response, higher inocula highly activate EphA2, leading to a strong inflammatory response.

To determine the biological significance of EphA2 as a β-glucan receptor for *C. albicans in vivo*, we tested the effects of DAS in the corticosteroid-treated mouse model of OPC (*23*). Mice that received DAS had increased severity of infection, as manifested by greater weight loss and higher oral fungal burden relative to the animals that received the vehicle control (Fig. 4A, B). Histopathological analysis of the tongues of DAS-treated mice confirmed that inhibition of EphA2 signaling resulted in larger fungal lesions and deeper tissue invasion (Fig. S19). Mice treated with DAS also had greater fungal dissemination to the liver, suggesting that EphA2 is required for maintaining the epithelial barrier function of the GI tract (Fig. 4C). Consistent with our *in vitro* results, we found that *C. albicans* infection of the vehicle control mice induced strong Stat3 phosphorylation in the oral epithelium, and this phosphorylation was reduced by treatment with DAS (Fig. S20). Administration of DAS also reduced *Il17a* mRNA expression in the oral tissues by 350-fold, *Il22* mRNA by 1000-fold, *S100a8* mRNA by 9-fold and *Defb3* mRNA by 7-fold, suggesting that EphA2 signaling is necessary for a maximal IL-17/IL-22 response (Fig. 4D). Thus, EphA2 is a key regulator of the host inflammatory response to *C. albicans* that limits fungal proliferation during oral infection.

**Fig. 4.**
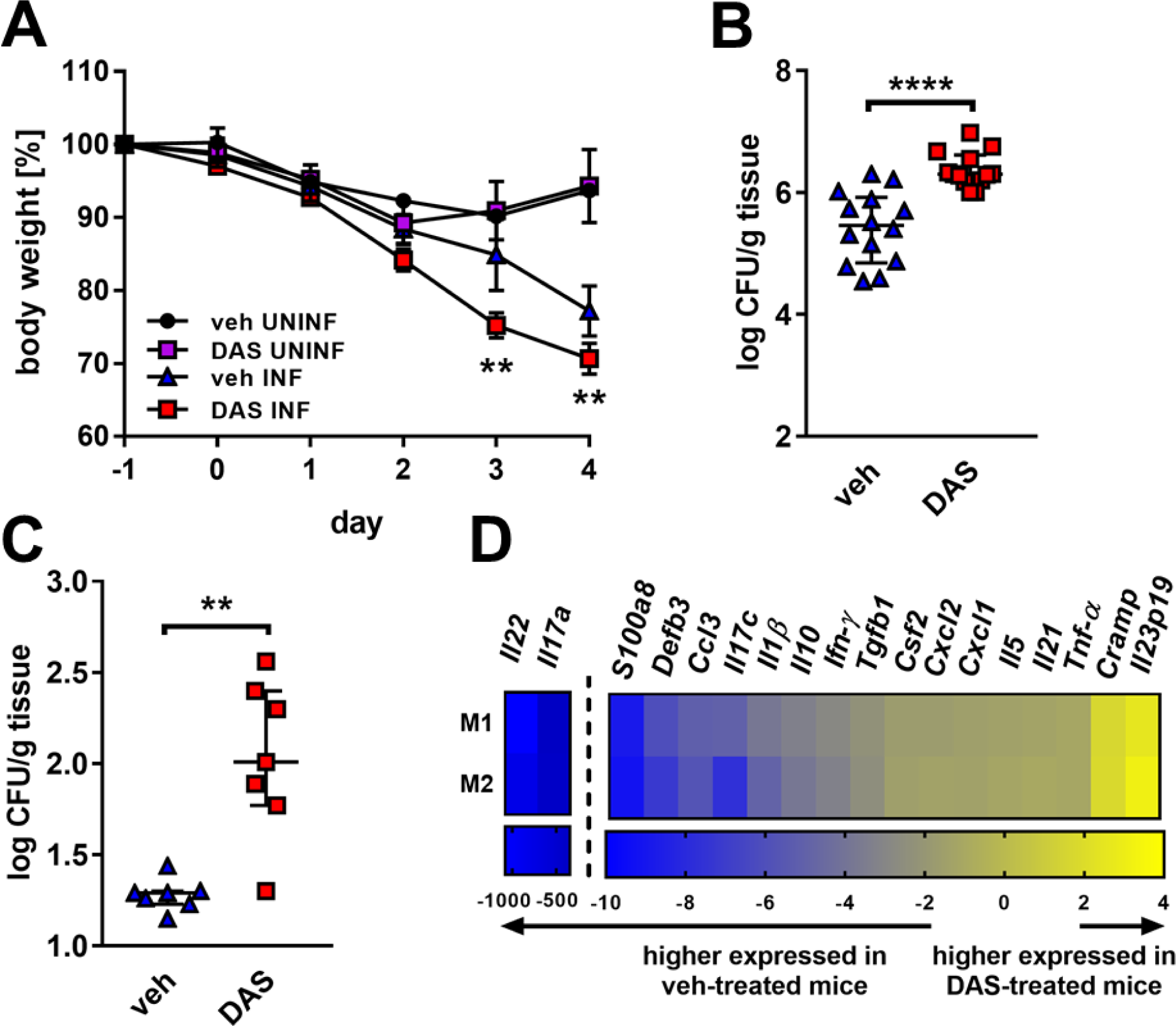
EphA2 signaling is required for resistance to OPC. (**A**) Effects of treatment with DAS on the body mass of mice with OPC. Results are the mean ± SD of 7 mice per group. (**B**) Oral fungal burden of mice with OPC after 4 d of infection. Results are the combined data from 2 independent experiments. Results are median ± interquartile range of 14 mice in the control group and 13 mice in the DAS group. (**C**) Liver fungal burden after 4 d of infection. Results are median ± interquartile range of 7 mice per group in a single experiment. (**D**) Effects of DAS on mRNA expression of inflammatory mediators in the mouse tongue after 4 d of infection. mRNA levels were normalized to GAPDH, and the results are presented as fold change relative to the vehicle control mice. Data from two mice (M1 and M2) are presented. Veh, vehicle. \**P* < 0.05, \*\**P* < 0.01, \*\*\**P* < 0.001, \*\*\*\**P* < 0.0001; NS, not significant.

The data presented herein indicate that EphA2 is a non-classical receptor for β-glucans that plays a vital role in sensing the presence of *C. albicans* yeast and hyphae in the oral cavity. While EphA2 contributes to receptor-induced endocytosis of the organism, this function is overshadowed by the central role of EphA2 in sensing the presence of high levels of fungal β-glucans, activating MAPKs and Stat3, and thereby stimulating epithelial cells to produce pro-inflammatory cytokines and HDPs. *In vivo*, β-glucan recognition of invading fungi via EphA2 is also required for the production of IL-17, IL-22, and β defensin 3, which are essential for resistance to OPC (*18, 24, 25*). Thus, our data support the model that EphA2 recognizes β-glucans expressed by pathogenic fungi and thereby primes epithelial cells to respond appropriately and prevent OPC when the fungal inoculum increases above threshold levels (Fig. S21).

## Acknowledgments

This work was supported in part by NIH grants R01DEO22600 and R01AI124566.

We thank Samuel W. French and Edward Vitocruz for histopathology, Ashraf S. Ibrahim and Michael R. Yeaman for providing strains, and members of the Division of Infectious Diseases at Harbor-UCLA Medical Center for critical suggestions.

S.G.F. is a cofounder of and shareholder in NovaDigm Therapeutics, Inc.

## Supplementary Materials

### Materials and Methods

#### Ethics statement

All animal work was approved by the Institutional Animal Care and Use Committee (IACUC) of the Los Angeles Biomedical Research Institute.

#### Fungal Strains, cell Lines, and reagents

The strains used in the experiments (listed in Table S2) were grown as described previously (*1*). For *C. albicans* interactions with OKF6/TERT-2 oral epithelial cell lines (provided by Jim Rheinwald, Dana-Farber/Harvard Cancer Center, Boston, MA) (*2*) were grown as described (*1, 2*). To determine the effects of the inhibitors, the host cells were treated 1 hour post infection with 2.5 μM dasatinib (*3*), 400 μM EphA2 antagonist, a 2,5-dimethylpyrrolyl benzoic acid derivative (*4, 5*), 50 μM Stat3 inhibitor S31-201 (*6*), 1μM gefitinib (*7*), 3 μg/ml dectin-1 mAb (R&D Systems) (*8*), 50 μg/ml depleted zymosan (InvivoGen), 50 μg/ml mannan (Sigma), or 50 μg/ml laminarin (InvivoGen). OKF6/TERT-2 cells were transfected with random control siRNA (Santa Cruz Biotechnology; sc-37007) or EphA2 siRNA (Santa Cruz Biotechnology; sc-29304) using Lipofectamine 2000 (Invitrogen) following the manufacturer’s instructions. Various experiments included the use of different Abs: EphA2 (Cell signaling; #6997; #6347), Stat3 (Cell signaling; #12640; #9134), c-Fos (Cell signaling; #5348; #4384), MEK1/2 (Cell signaling; #9122; #9154), p65 (Cell signaling; # 3033 # 8242), β-actin (Cell signaling # 3700), human dectin-1/CLEC7A (R&D Systems; #MAB1859), Fc-hDectin-1a (Invivogen), and Fc-mDectin-1a (Invivogen). Recombinant cytokines and proteins: IL-17 (Peprotech), TNF-α (Peprotech), EFNA1 (Acro Biosystems), and EphA2 (BioLegend).

As described previously (*7*), siRNA was used to deplete EphA2 from the epithelial cells. OKF6/TERT-2 cells were transfected with random control siRNA or EphA2 siRNA (80 pmol; Santa Cruz Biotechnology) using Lipofectamine 2000 (Thermo Fisher Scientific) following the manufacturer’s instructions.

#### Confocal Microscopy

The accumulation of EphA2 around *C. albicans* was visualized using a minor modification of our previously described methods (*1*). Oral epithelial cells were infected with 2 × 10^5^ yeast organims of a wild-type *C. albicans* strain expressing GFP. After 15, and 90 min, the cells were fixed in 3% paraformaldehyde (wt/vol), blocked with 10% BSA (vol/vol), and incubated with antibodies against total EphA2, Stat3, or *C. albicnas* (*9*) followed by the appropriate secondary antibodies that had been labeled with either AlexaFluor 488 or AlexaFluor 568. The cells, or OCT sections, were then imaged by confocal microscopy, and the final images were generated by stacking optical sections along the *z* axis.

#### Flow Cytometry

Human buccal epithelial cells were isolated from healthy donors by scraping the oral cavity. The cells were suspended in DMEM + 10% FBS Pen/Strep for 1 hours followed by washing in HBSS. Cells were incubated for 1 hour with Dectin-1 mAB (R&D) followed by anti-mouse AlexaFluor 488 Ab. Control cells were incubated in a similar concentration of mouse IgG (Abcam, Inc.). The fluorescence of the cells was determined by flow cytometry, analyzing at least 10,000 cells per condition.

#### Detection of Phosphorylation

OKF6/TERT-2 cells in 24-well tissue culture plates were infected with 1 × 10^6^ *C. albicans* for various times. Next, the cells were rinsed with cold HBSS containing protease and phosphatase inhibitors and removed from the plate with a cell scraper. The cells were collected by centrifugation and boiled in sample buffer. The lysates were separated by SDS/PAGE, and the phosphorylated proteins were detected by immunoblotting with phospho-specific antibodies. The blots were then stripped, and total protein levels and β-actin were detected by immunoblotting.

#### Stat3 ELISA

To quantify the phosphorylation of Stat3, OKF6/TERT-2 cells in 96-well tissue culture plates were treated with inhibitors for 1 h and then infected with 3 × 10^5^ *C. albicans* cells for different time periods. Next the epithelial cells were permeabilized and the phosphorylation of Stat3 was measured by an ELISA for phosph-Ser727 (LSBio) following the manufacturer’s instructions.

#### Measurement of Host Cell Endocytosis

The endocytosis of *C. albicans* by oral epithelial cells was quantified as described previously (*1*). OKF6/TERT-2 cells were grown to confluency on fibronectin-coated circular glass coverslips in 24-well tissue culture plates. They were infected with either 2 × 10^5^ yeast-phase *C. albicans* cells per well for 120 min. To determine the effects of the inhibitors on endocytosis, the host cells were incubated with indicated inhibitors as described above. Control cells were incubated in a similar concentration of diluent (0.25% DMSO). The inhibitors were added to the host cells 60 min before the fungal cells, and they remained in the medium for the entire incubation period.

#### Host cell damage assay

The extent of host cell damage caused by the *C. albicans* was measured using a standard ^51^Cr release assay (*1, 10*). Briefly, OKF6/TERT-2 were grown to confluence in 96-well tissue culture plates, loaded with ^51^Cr overnight, and then infected with 3 × 10^5^ cells of the *C. albicans*. Parallel wells were left uninfected as a negative control. After an 8-h incubation, the amount of ^51^Cr that had been released into the medium and that remained associated with the cells was measured. To determine the effects of the inhibitors on cell damage, the host cells were incubated with 2.5 μM dasatinib or 400 μM EphA2 antagonist. Control cells were incubated in a similar concentration of diluent (0.25% DMSO). Each experiment was performed three times in triplicate.

#### RNA isolation and RT-PCR

Total RNA was isolated and RT-PCR was performed as previously described (*11*). For RT-PCR host RNA was extracted using the Ribopure Yeast Kit (Life Technologies), according to the manufacturer’s instructions. The mRNA levels of were measured by real-time PCR using the primers listed in Table S3 in the supplemental material. The relative transcript level of each gene was normalized to GAPDH by the 2^-ΔΔCT^ method.

#### Measurement of hBD2 expression

To determine hBD2 expression in OKF6/TERT-2 cells peptide secretion was performed following the manufacturer’s instructions. Briefly, OKF6/TERT-2 cells in 24-well tissue culture plates were pre-treatment of inhibitors following infection with 1 × 10^6^ *C. albicans* for 20 hours. The supernatant was collected and hBD2 secretion was determined by ELISA.

#### IL-8 Elisa

To determine IL-8 release of OKF6/TERT-2 cells supernatants were collected and ELISA was determined following manufators instructions (R&D Systems).

#### Cytokine array

Cytokine levels in culture supernatants were determine as previously described (*12*). Briefly OKF6/TERT-2 cells were infected with *C. albicans* in a 96-well plate (MOI 5). After 8 h infection, the supernatant was collected, clarified by centrifugation and stored in aliquots at -80 °C. The concentration of IL-1α, IL-1β, and IL-8/CXCL8 in the medium was determined using the Luminex multipex assay (R&D Systems). Each condition was tested in three independent experiments.

#### EphA2-binding ELISA

50 μg carbohydrates (laminarin, zymosan, or mannan), 2 μg EFNA1 (positive control), or 50 μg BSA were coated ON at 20 °C in 96-well plates, blocked with non-fat dry milk for 1 hour. After washing 100 μg/mL recombinant EphA2 (Gln25-Asn534-6xHis; BioLegend) was added to each well and incubated for 2 hours at 37 °C. Washing was repeated and 1 μg/mL anti-His-HRP mAb was added and incubated for 1 hour at 37 °C. Washing was repeated and color reagent was added following the manufacturer’s instructions (R&D Systems). After 5 min stop solution was added and values were read at 450 nm.

#### Mouse model of oropharyngeal candidiasis

The effect of DAS on the severity of OPC was determined using a mouse model of oropharyngeal candidiasis (*13*). Male BALB/C mice were fed an oral solution of dasatinib (10mg/kg/day), twice a day, administered at 0.05 ml/ dose starting on day -1 relative to infection. Triamcinolone (15 mg/kg) was administered s.c. on days -1, 1, and 3 (*13*). For inoculation, the mice were sedated, and a swab saturated with 10^6^ *C. albicans* cells was placed sublingually for 75 min; 4 d later, the mice were euthanized, and their tongue and attached tissues were harvested and divided by two. One-half was weighed, homogenized, and quantitatively cultured. The other one-half was snap frozen in Optimal Cutting Temperature (OCT); 2 μm-thick sections were cut with a cryostat and fixed with cold acetone. To detect phosphorylation of Stat3, the cryosections were rehydrated in PBS and then blocked with 10 % BSA. Sections were stained with P-Stat3 Ab (Cell Signaling), primary antibodies rinsed, and stained with an AlexaFluor 488 secondary antibody followed by anti-*Candida* AlexaFluor 568 and imaged by confocal microscopy. To enable comparison of fluorescence intensity between slides, the same image acquisition settings were used for each experiment. For histopathology, additional cryosections were prepared and stained with periodic acid-Schiff (PAS) stain.

#### Statistics

At least three biological replicates were performed for all *in vitro* experiments unless otherwise indicated. Data were compared by Mann-Whitney or unpaired Student's *t* test using GraphPad Prism (v. 6) software. *P* values < 0.05 were considered statistical significant.

---

**Table S1.**
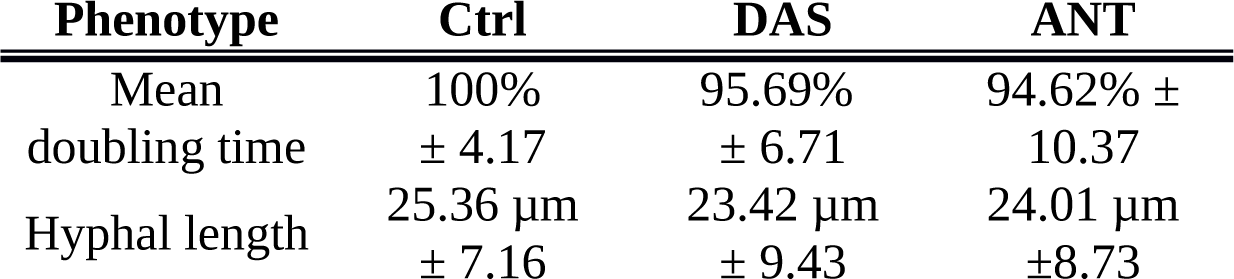
Effect of EphA2 inhibitors on fungal growth

**Table S2.** Strains used in this study

| Strain | Reference |
| --- | --- |
| *C. albicans* SC5314 | (1) |
| *C. albicans* CAI4-GFP | (2) |
| *C. albicans* 529L | (3) |
| *C. albicans* *efg1/cph1* | (4) |
| *C. glabrata* 05-761 | (5) |
| *S. cerevisiae* BY4743 | (6) |
| *R. delemar* 99-880 | (7) |
| *A. fumigatus* Af293 | (8) |
| *E. coli* K12 | (9) |
| *S. aureus* SH1000 | (10) |

**Table S3.**
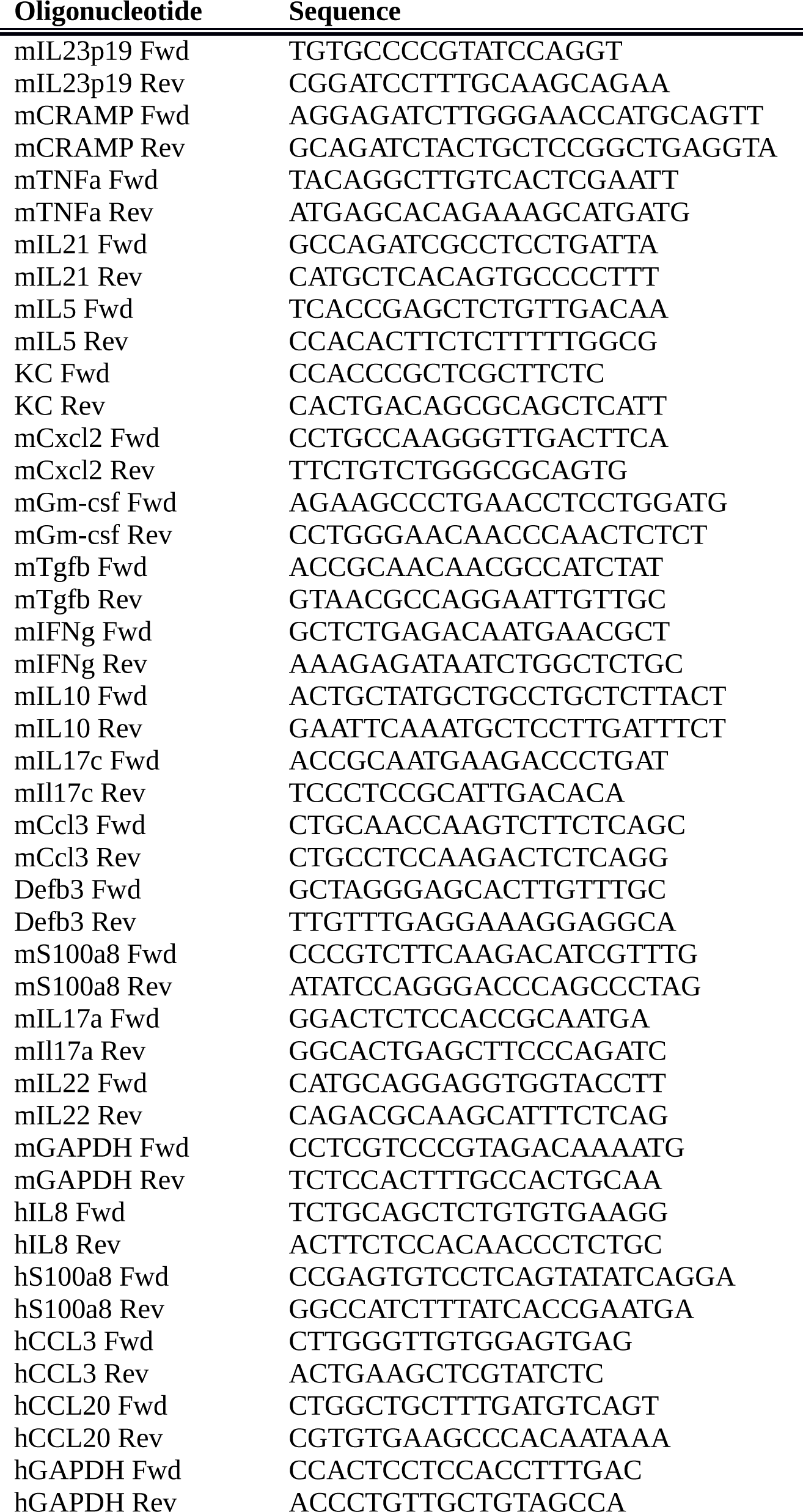
Oligonucleotides for RT-PCR

**Fig. S1.**
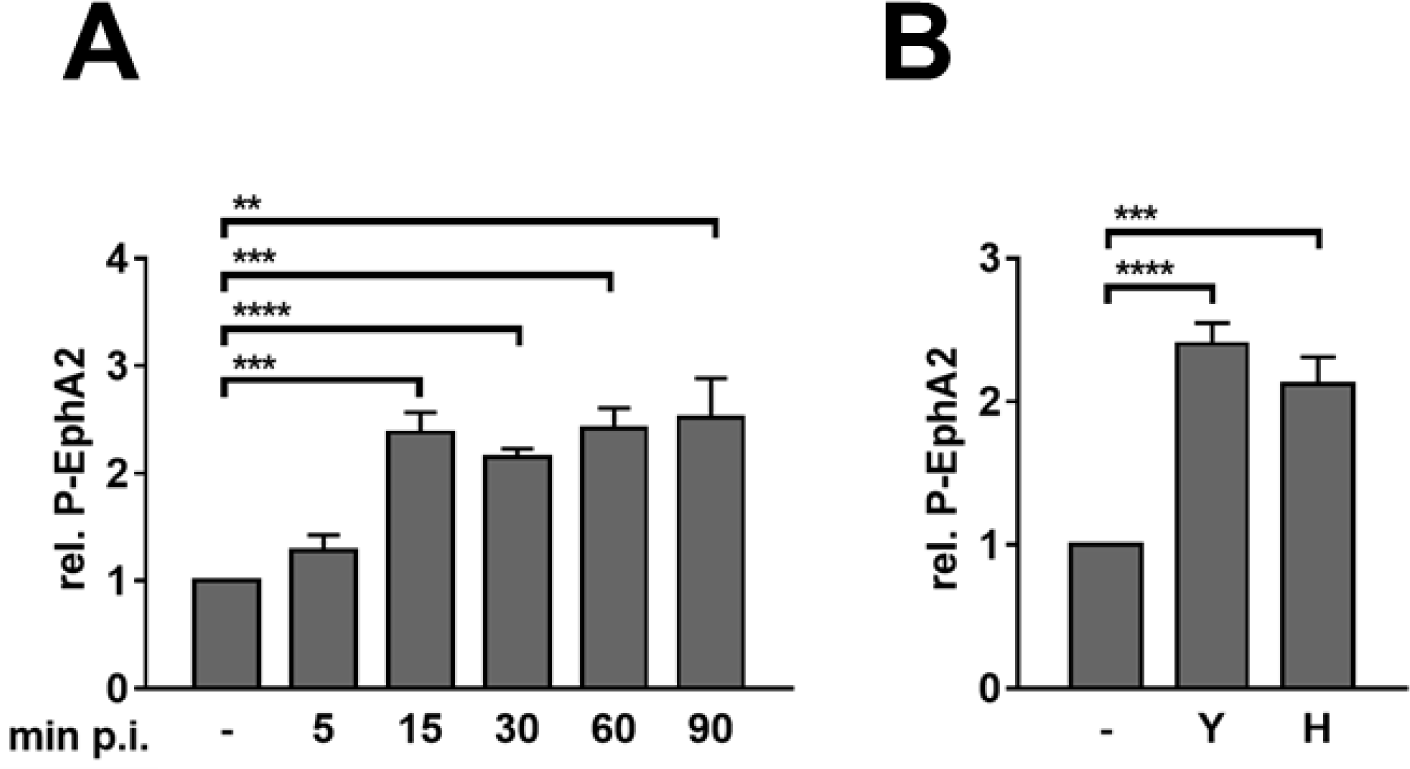
Densitometric analysis of EphA2 phosphorylation during *C. albicans* infection of oral epithelial cells. (**A**) EphA2 phosphorylation in oral epithelial cells infected with yeast-phase *C. albicans* SC5314 for the indicated times. (**B**) EphA2 phosphorylation in epithelial cells exposed to either yeast-phase or hyphal-phase *C. albicans* for 20 min. Graphs show the relative ratio of phosphorylated EphA2 to total EphA2. Data are the mean ± SD of 3 independent immunoblots. H, hyphae; p.i., post-infection; Y, yeast. \*\**P* < 0.01, \*\*\**P* < 0.001, \*\*\*\**P* < 0.0001.

**Fig. S2.**
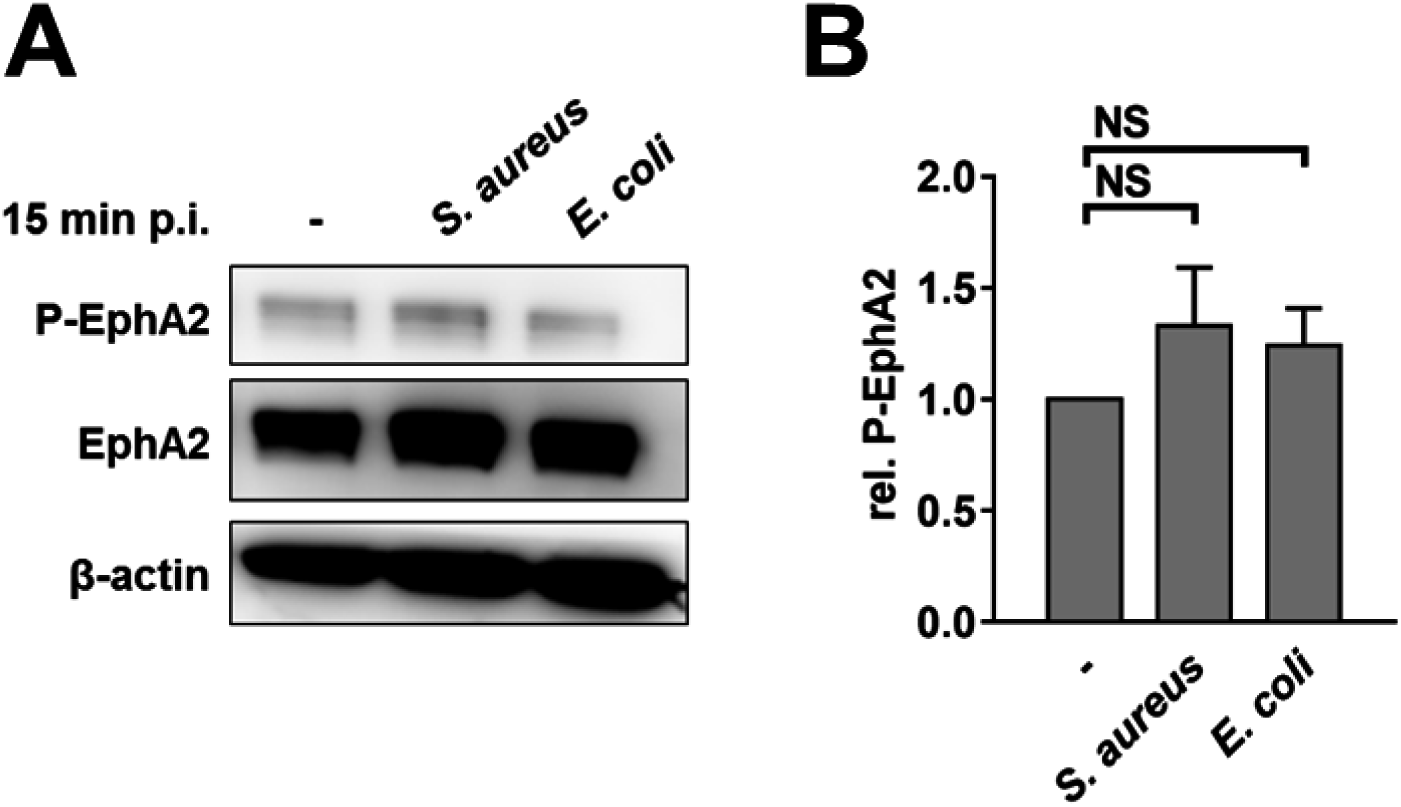
EphA2 is not activated by epithelial cell infection with *Staphylococcus aureus* or *Escherichia coli*. (**A**) Immunoblot showing EphA2 phosphorylation of epithelial cells after 15 min infect infection *S. aureus* SH1000 and *E. coli* K12 at multiplicity of infection of 5 organisms per epithelial cells. (**B**) Densitometric analysis of immunoblots as in (**A**). Data are the mean ± SD of 3 independent immunoblots. NS, not significant.

**Fig. S3.**
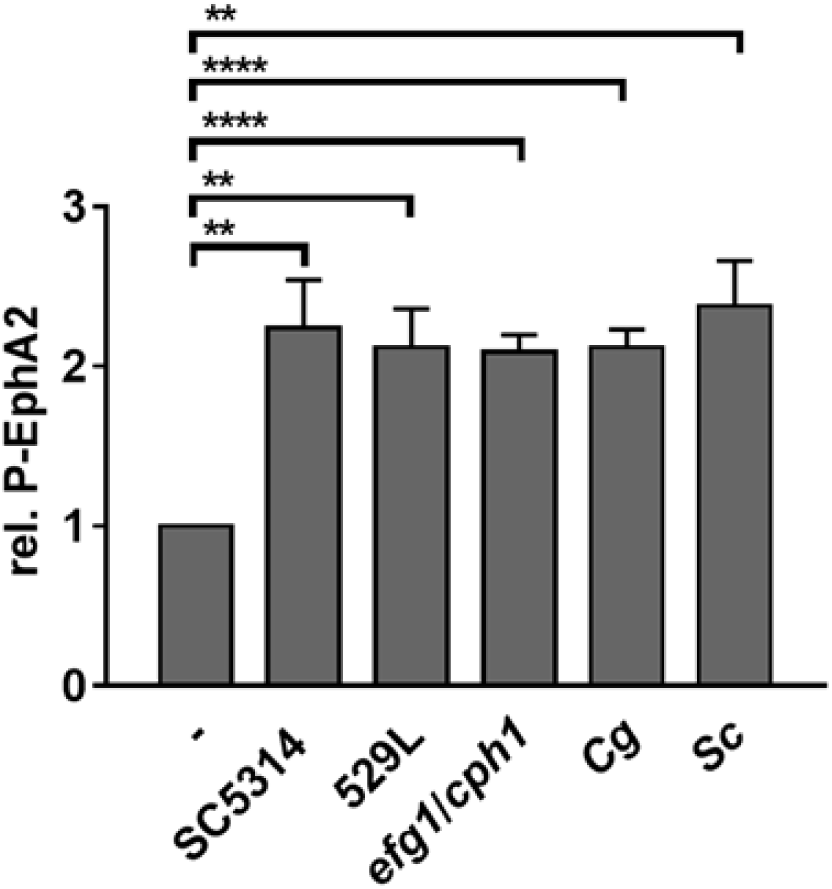
Effects of *C. albicans* (SC5314, 529L, *efg1/cph1*), *Candida glabrata*, and *Saccharomyces cerevisiae* on EphA2 phosphorylation. Cg, *C. glabrata*; Sc, *S. cerevisiae*. Data are the mean ± SD of 3 independent immunoblots. \*\**P* < 0.01, \*\*\*\**P* < 0.0001.

**Fig. S4.**
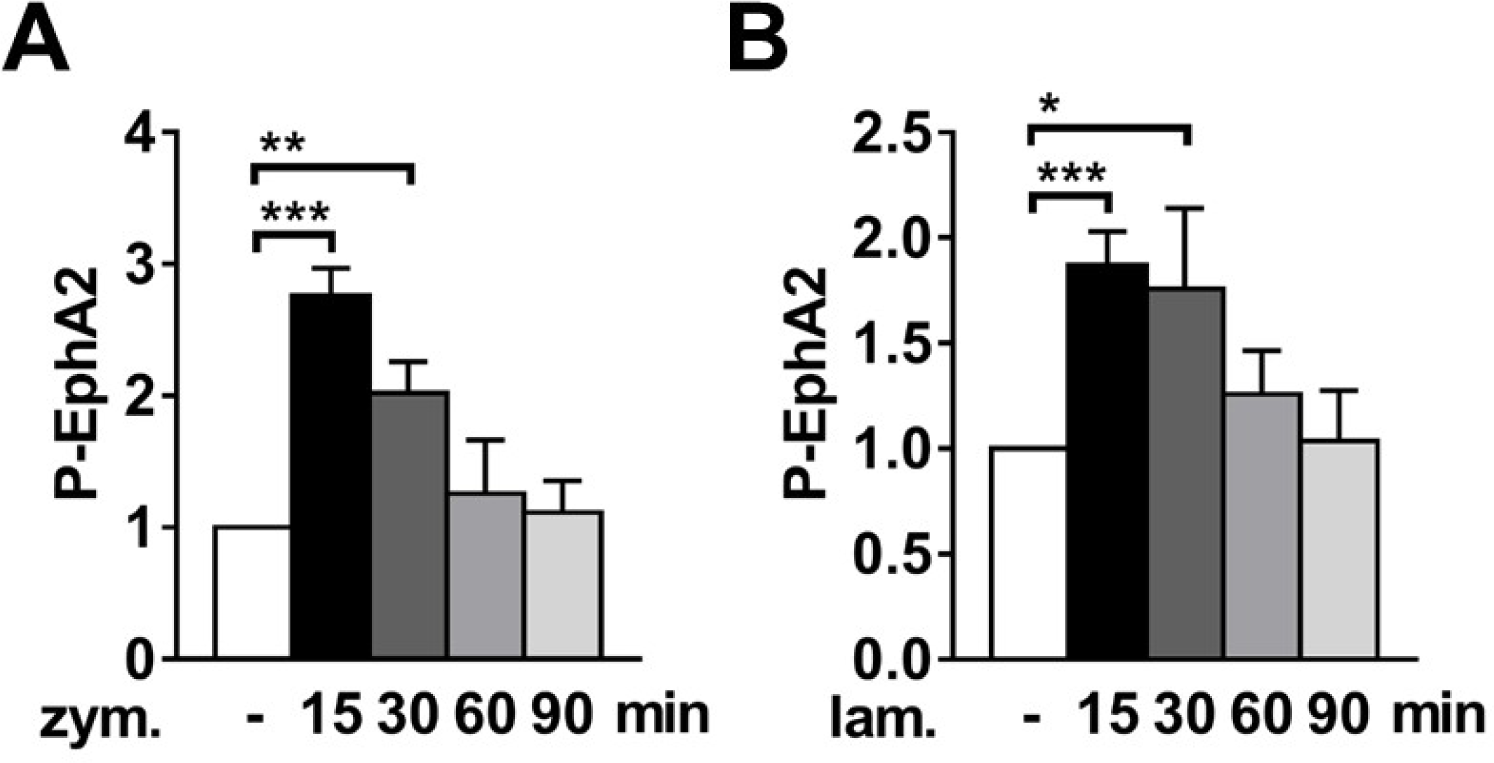
Densitometric analysis of EphA2 phosphorylation induced by the β-glucans zymosan and laminarin. EphA2 phosphorylation in epithelial cells that were incubated for the indicated times with (**A**) zymosan (50 μg/ml) and (**B**) laminarin (50 μg/ml). Data are the mean ± SD of 3 independent immunoblots. Lam, laminarin; zym, zymosan. \**P* < 0.05, \*\*\**P* < 0.001.

**Fig. S5.**
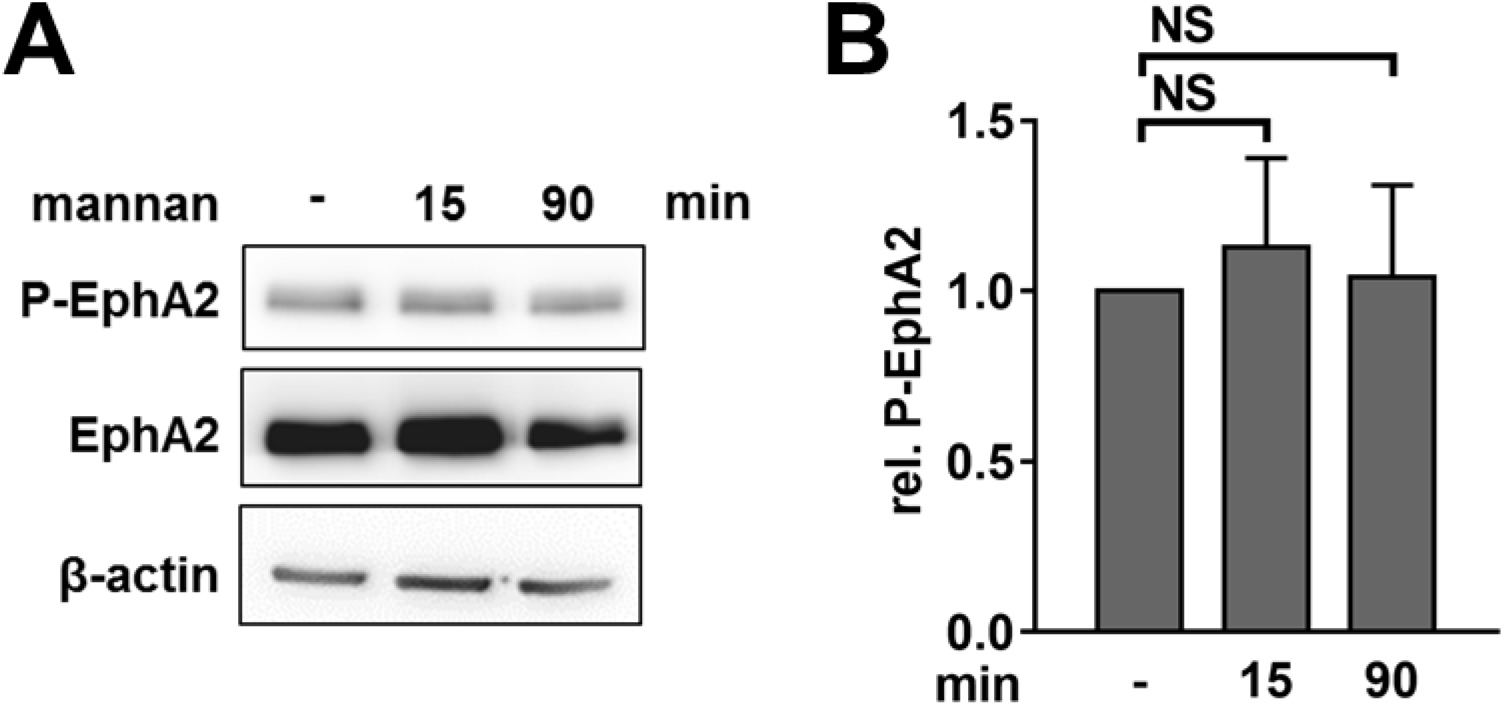
Mannan does not activate EphA2. Extent of EphA2 phosphorylation in epithelial cells that were incubated for 15 and 90 min with mannan (50μg/mL). (**A**) Representative immunoblot. (**B**) Densitometric analysis. Data are the mean ± SD of 3 independent immunoblots. NS, not significant.

**Fig. S6.**
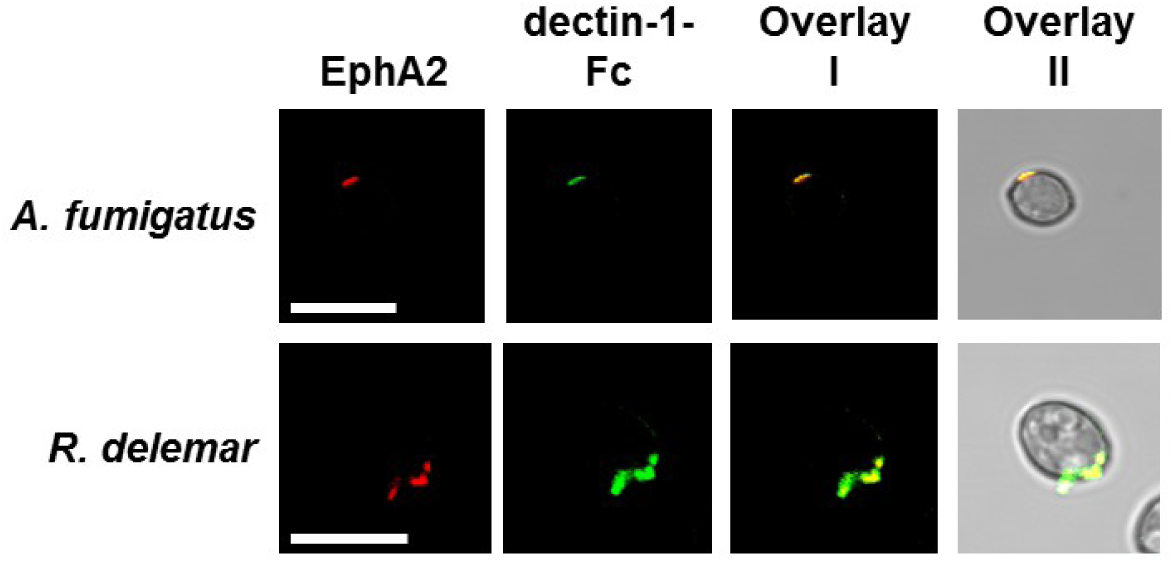
EphA2 and dectin-1-Fc co-localize on the surface of swollen spores and conidia of filamentous fungi. Images of swollen conidia of *Aspergillus fumigatus* and *Rhizopus delemar* that were stained with recombinant EphA2 (red) and dectin-1-Fc (green). Scale bar 10 μm.

**Fig. S7.**
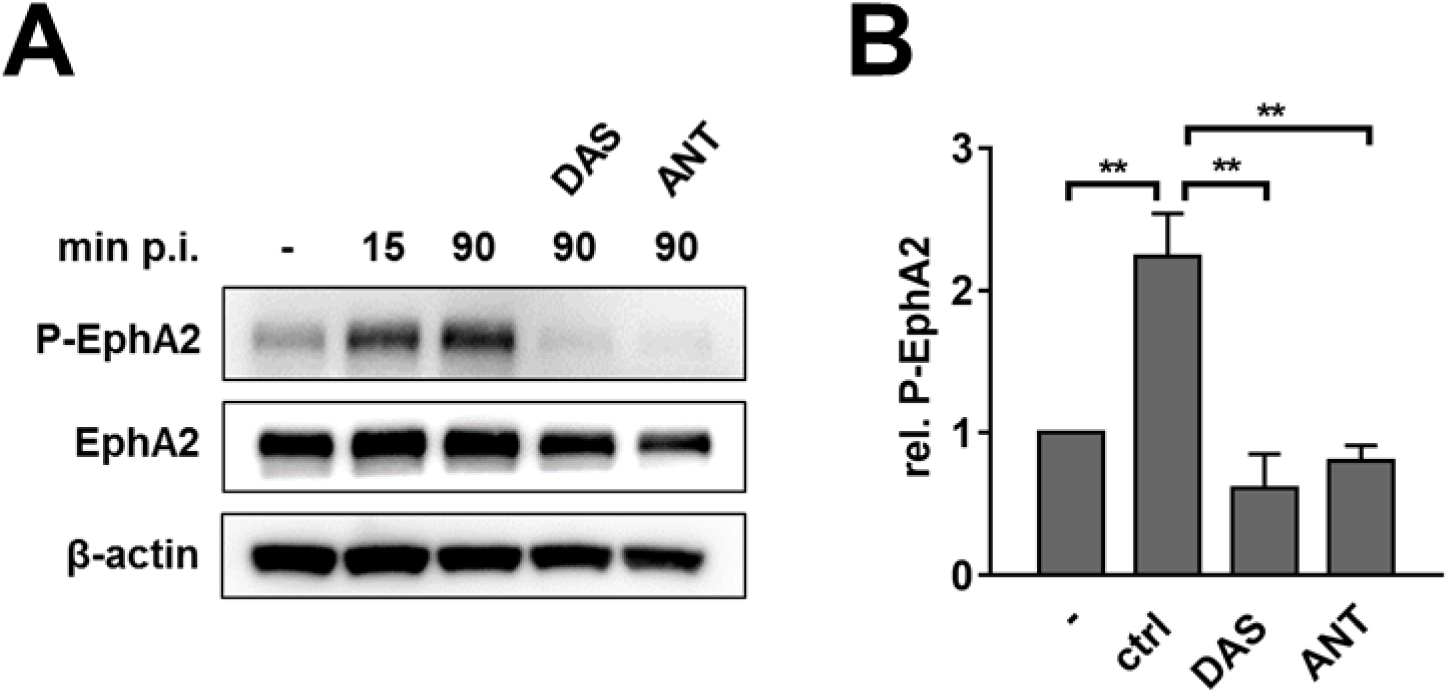
EphA2 inhibitors block *C. albicans*-induced EphA2 activation. (**A**) EphA2 phosphorylation in oral epithelial cells that were preincubated with dasatinib or an EphA2 antagonist and then infected with *C. albicans*. (**A**) Representative immunoblot. (**B**) Densitometric analysis. Results are the mean ± SD of 3 independent immunoblots. Ctrl, control; DAS, dasatinib; ANT, EphA2 antagonist. \*\**P* < 0.01.

**Fig. S8.**
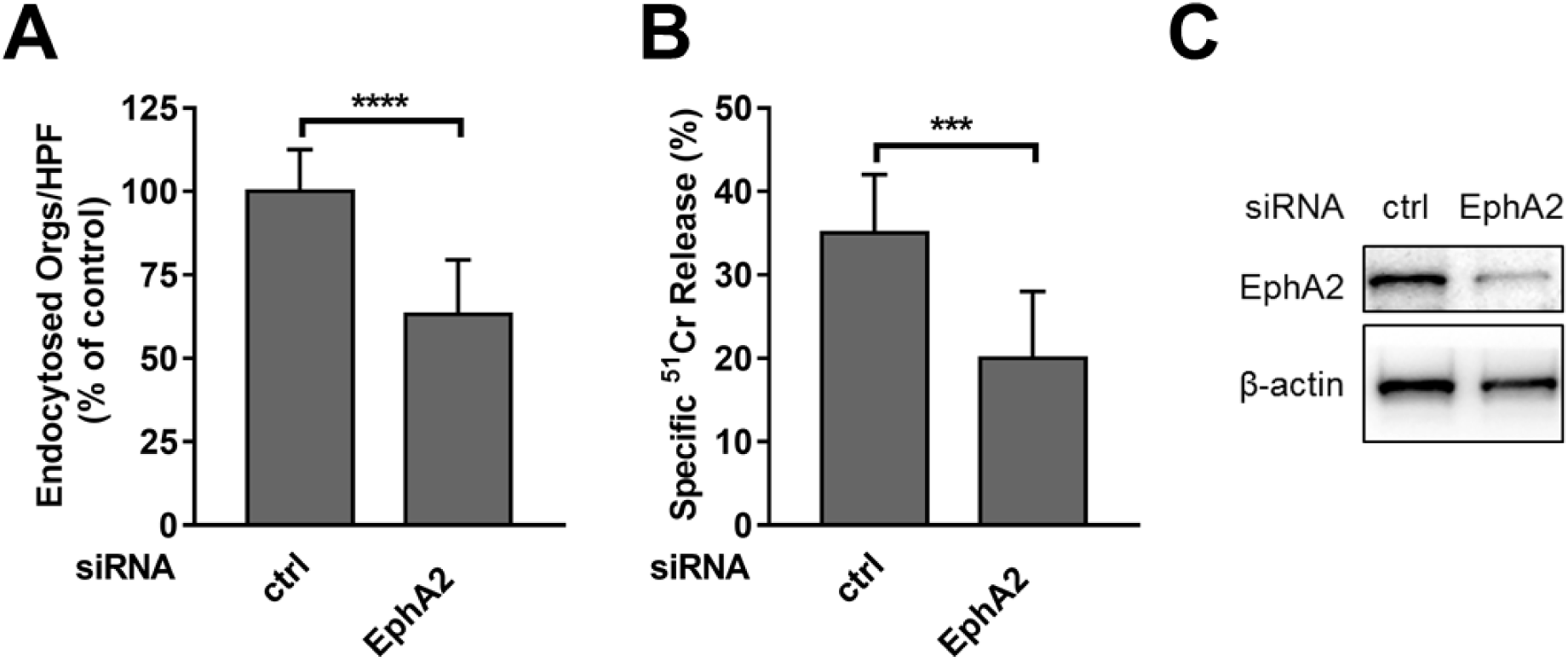
Depletion of EphA2 with siRNA in oral epithelial cells reduces *C. albicans* endocytosis and cell damage. Oral epithelial cells were transfected with either random control siRNA or EphA2 siRNA and then infected with *C. albicans*. Effects of siRNA on (**A**) epithelial cell endocytosis and (**B**) *C. albicans-*induced epithelial cell damage. Data are the mean ± SD of 3 independent experiments, each performed in triplicate. (**C**) Representative immunoblot showing EphA2 depletion using siRNA. Ctrl, control. \*\*\**P* < 0.001, \*\*\*\**P* < 0.0001.

**Fig. S9.**
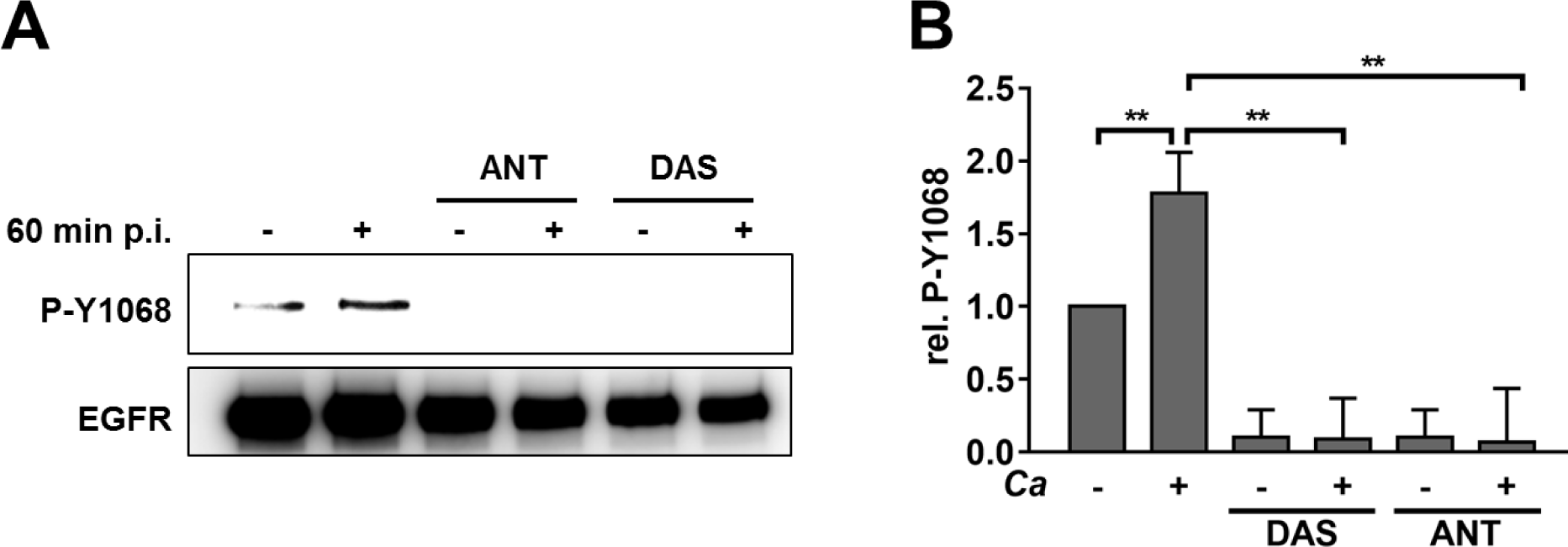
EphA2 inhibition decreases *C. albicans*-induced phosphorylation of the epidermal growth factor receptor (EGFR). EGFR phosphorylation at Y1068 in oral epithelial cells that had been pre-treated for 1 h with EphA2 inhibitors (DAS and ANT) and then infected for 1 h with *C. albicans*. (**A**) Representative immunoblot. (**B**) Densitometric analysis. Data are the mean ± SD of 3 independent immunoblots. \*\**P* < 0.01.

**Fig. S10.**
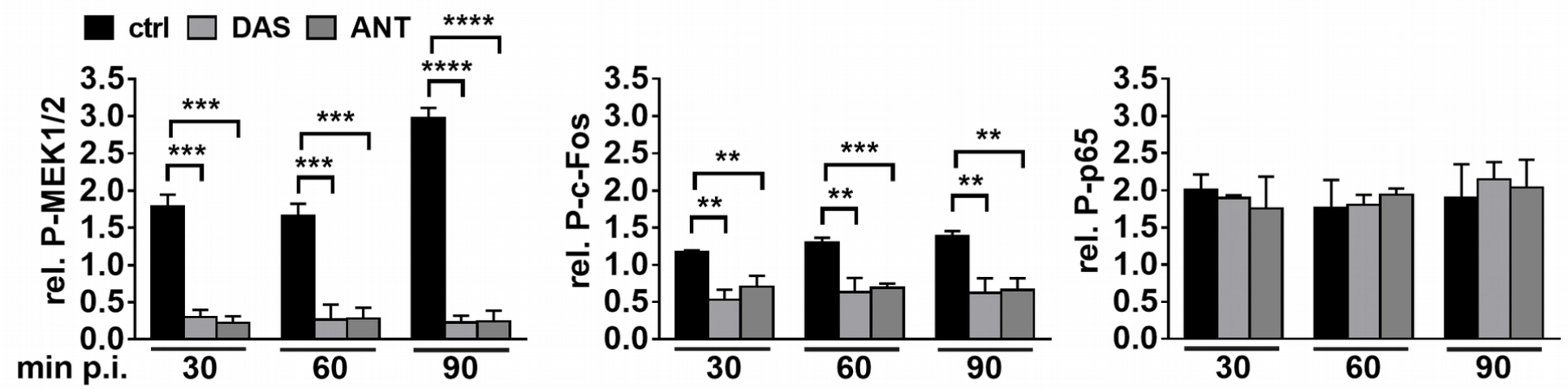
Effects of EphA2 inhibition on phosphorylation of MEK1/2, c-Fos, and p65. Data in each panel are the mean ± SD of the densitometric analysis of 3 independent immunoblots. \*\**P* < 0.01, \*\*\**P* < 0.001, \*\*\*\**P* < 0.0001.

**Fig. S11.**
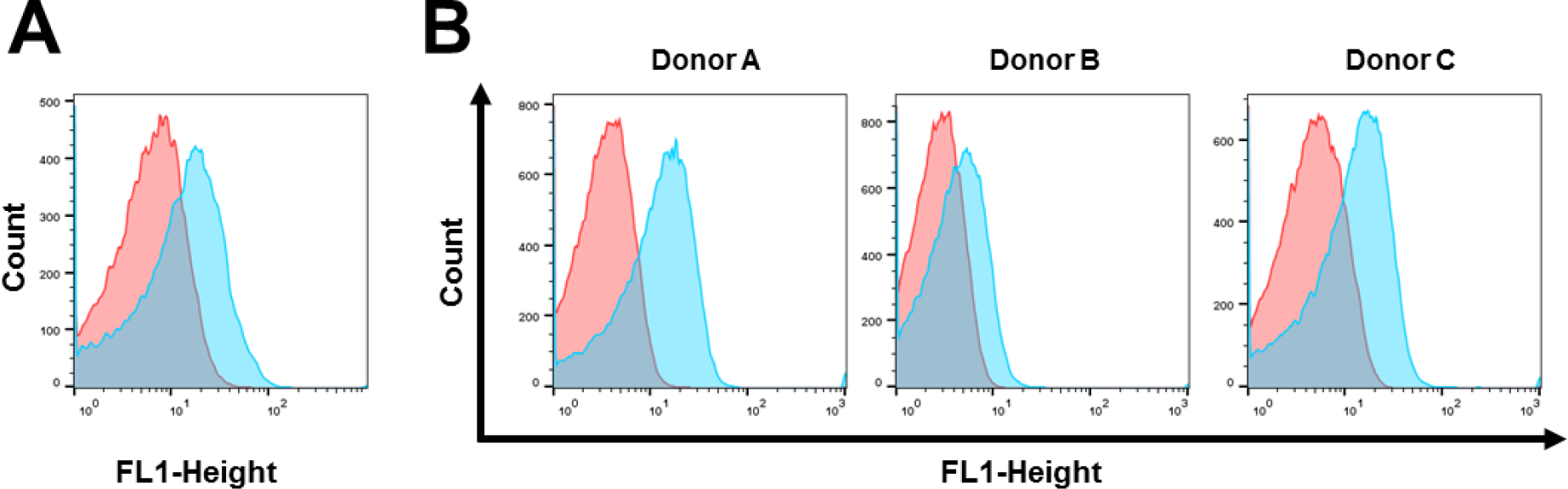
Dectin-1 is expressed on the surface of human oral epithelial cells. Flow cytometric analysis of dectin-1 surface expression on (**A**) the OKF6/TERT-2 human epithelial cell line and (**B**) human buccal epithelial cells from 3 different donors. Results from OKF6/TERT-2 or human buccal epithelial cells stained with the control IgG are shown in red and results from cells stained with dectin-1 antibody are shown in blue.

**Fig. S12.**
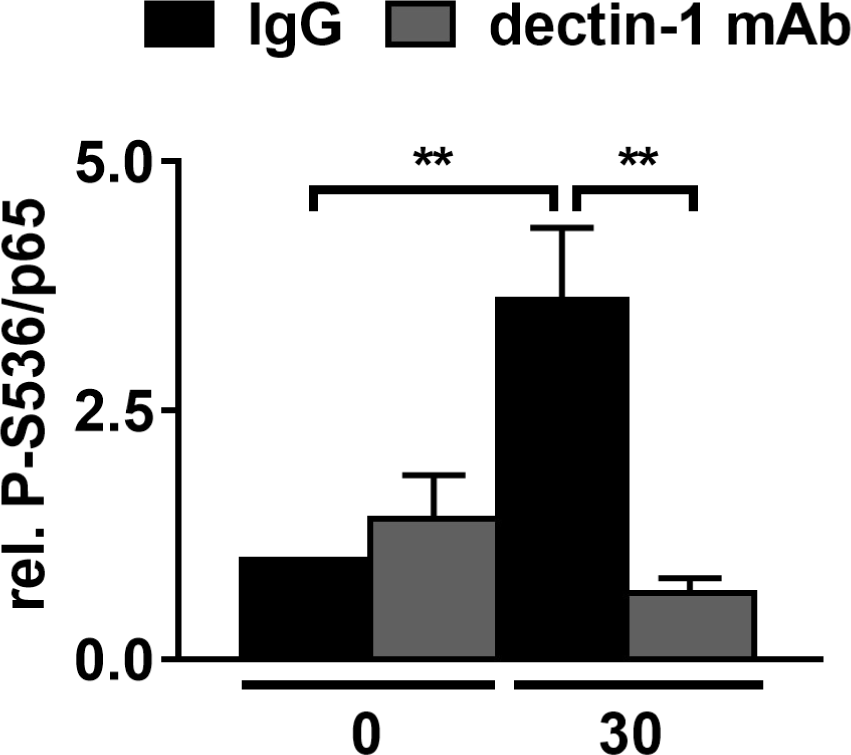
Densitometric analysis of NF-κB p65 phosphorylation. Oral epithelial cells were pretreated with control IgG or a dectin-1 mAb, and then infected with *C. albicans*. Data are the mean ± SD of 3 independent immunoblots.\*\**P* < 0.01.

**Fig. S13.**
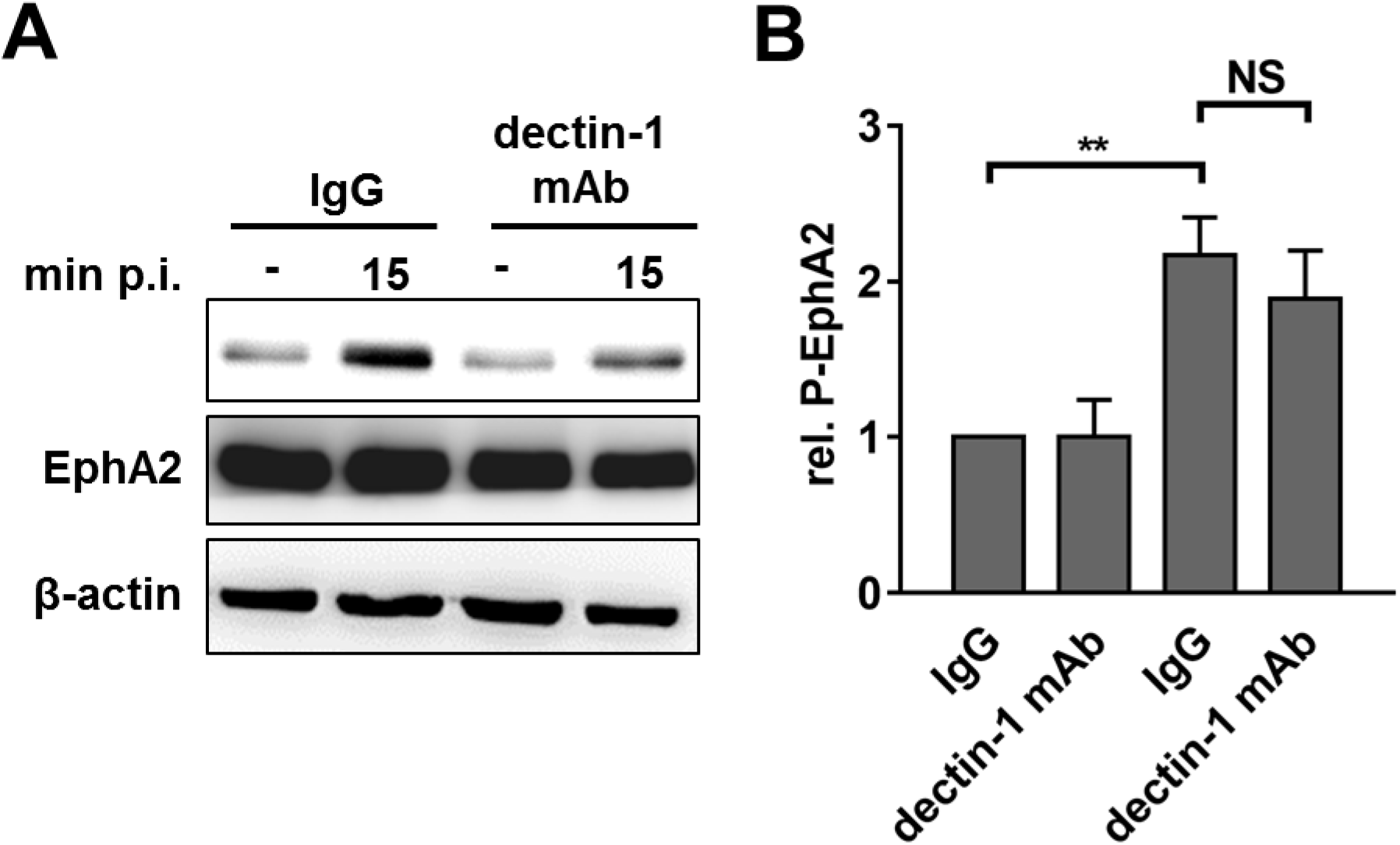
EphA2 phosphorylation is independent of dectin-1 signaling in human oral epithelial cells. (**A**) Representative immunoblot showing EphA2 phosphorylation in epithelial cells that were pre-treated with control IgG or a dectin-1 neutralizing antibody for 1 h prior to *C. albicans* infection. (**B**) Densitometric analysis of immunoblots such as the one in (**A**). Data are the mean ± SD of 3 independent immunoblots. \*\**P* < 0.01; NS, not significant.

**Fig. S14.**
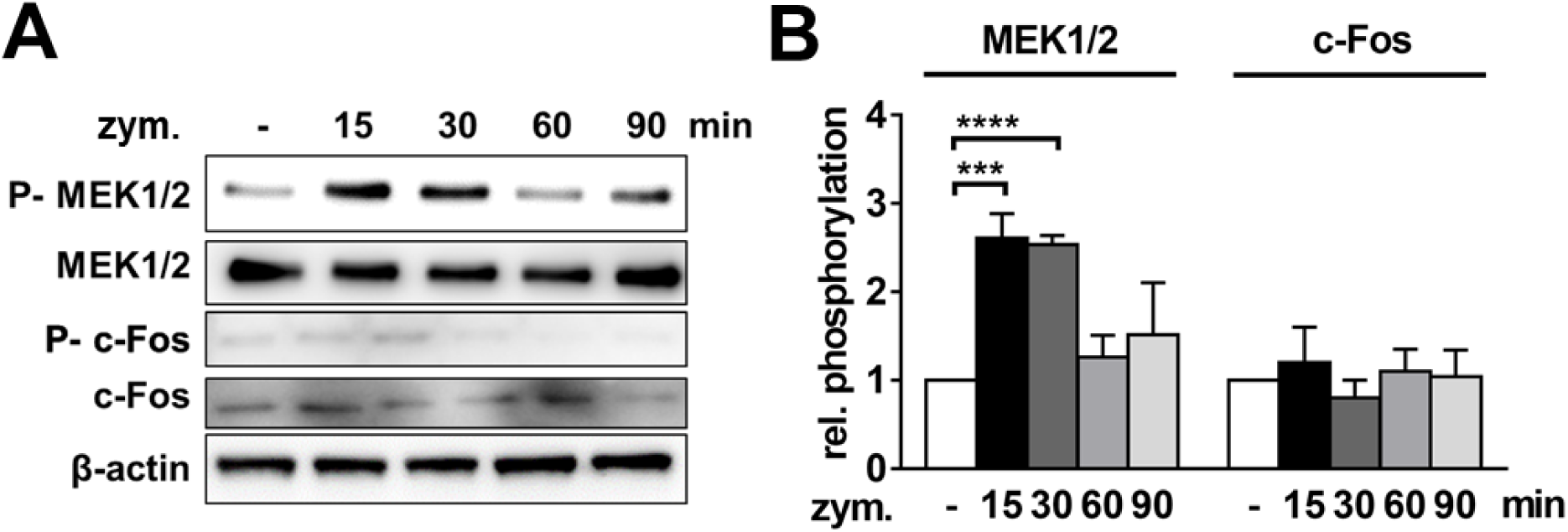
Zymosan induces transient MEK1/2 activation. (**A**) Representative immunoblot showing the phosphorylation of EphA2 and c-Fos in oral epithelial cells that had been incubated with zymosan (50 μg/mL) for the indicated time. (**B**) Densitometric analysis of immunoblots such as the ones in (**A**). Data are the mean ± SD of 3 independent immunoblots.\*\*\**P* < 0.001, \*\*\*\**P* < 0.0001.

**Fig. S15.**
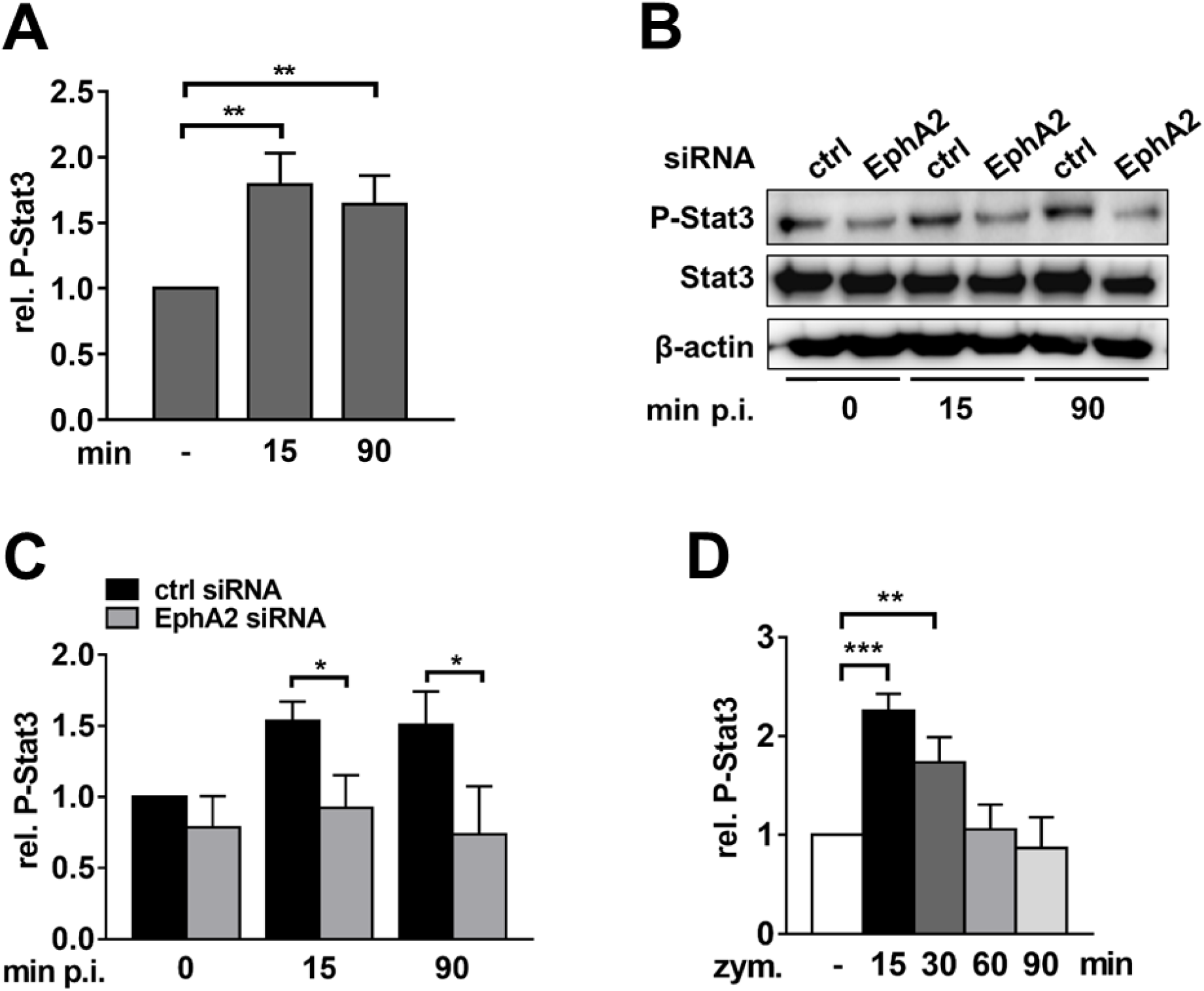
Phosphorylation of Stat3. (**A**) Stat3 is activated during *C. albicans* infection. Densitometric analysis of immunoblots showing *C. albicans*-induced Stat3 phosphorylation. Data are the mean ± SD of 3 independent immunoblots. (**B-C**) Effect of siRNA depletion of EphA2 on Stat3 phosphorylation in epithelial cells infected with *C. albicans*. (**B)** Representative immunoblot. (**C**) Densitometric analysis of immunoblots such as the one in (**B**). Data are the mean ± SD of 3 independent immunoblots. (**D**) Densitometric analysis of zymosan-induced Stat3 phosphorylation. Data are the mean ± SD of 3 independent immunoblots. \**P* < 0.05, \*\**P* < 0.01, \*\*\**P* < 0.001.

**Fig. S16.**
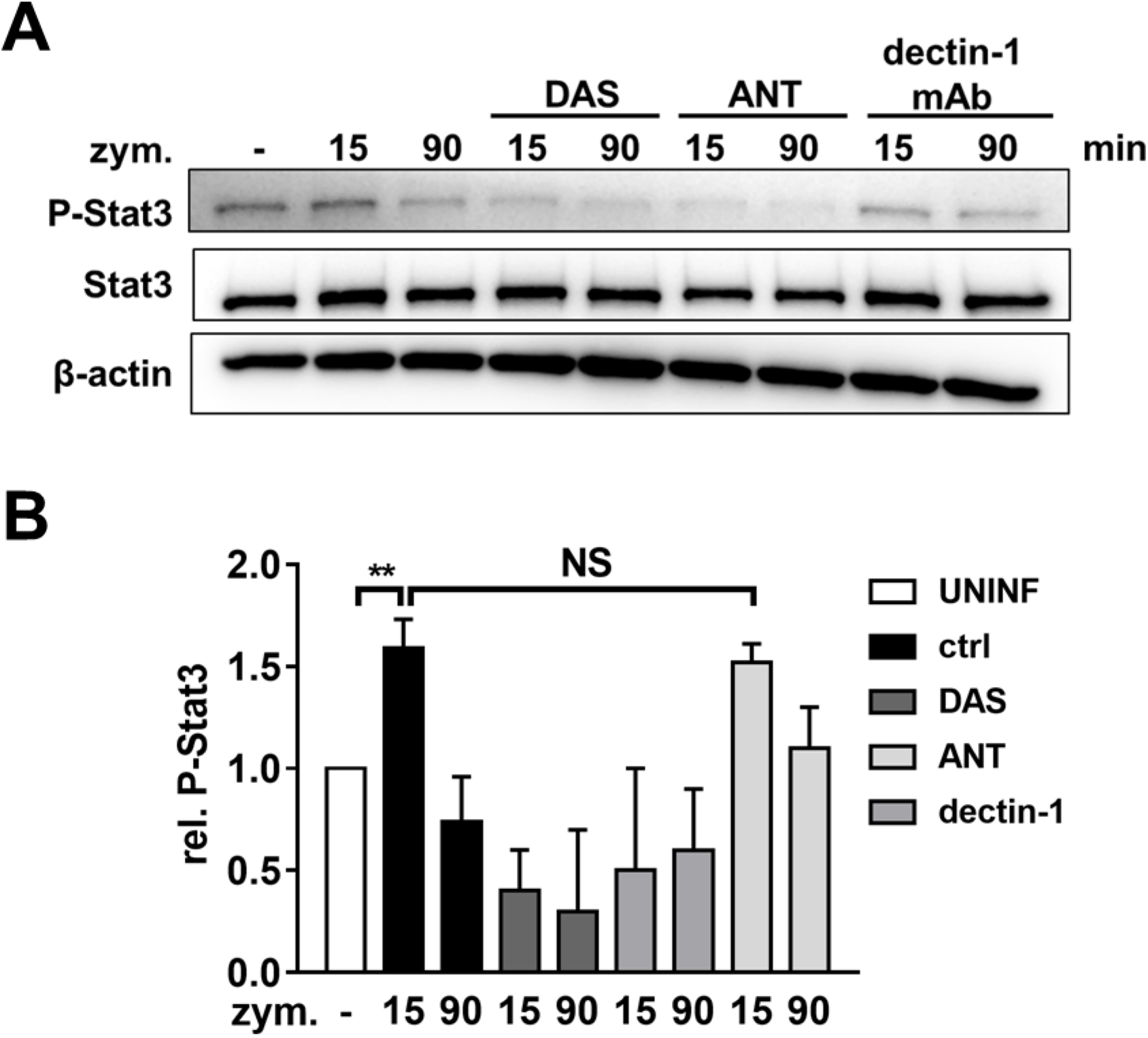
Stat3 activation requires EphA2, but not dectin-1. Stat3 phosphorylation in oral epithelial cells were incubated for 1 h with DAS, ANT, or a neutralizing anti-dectin-1 mAb, and then exposed to zymosan for indicated times. (**A**) Representative immunoblot (**B**) Densitometric analysis. Data are the mean ± SD of 3 independent immunoblots. \*\**P* < 0.01; NS; not significant.

**Fig. S17.**
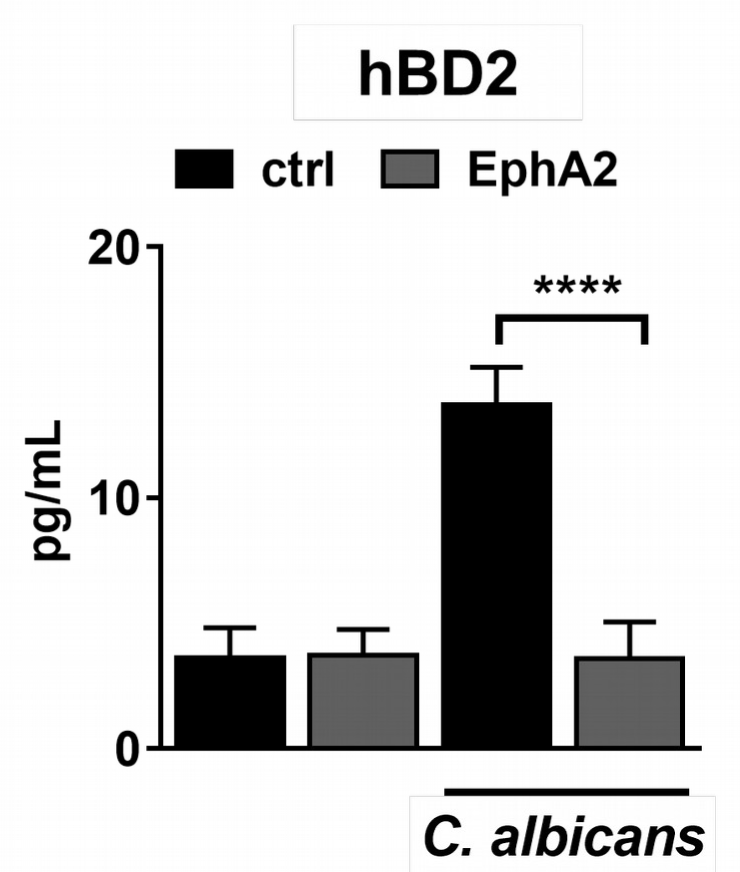
EphA2 depletion reduces hDB2 secretion in response to *C. albicans* infection. Human β defensin 2 secretion by oral epithelial cells that were transfected with either control siRNA or EphA2 siRNA and the infected for 8 h with *C. albicans*. Results are the mean ± SD of 3 independent experiments, each performed in duplicate. hBD2, human β defensin 2. \*\*\**P* < 0.001.

**Fig. S16.**
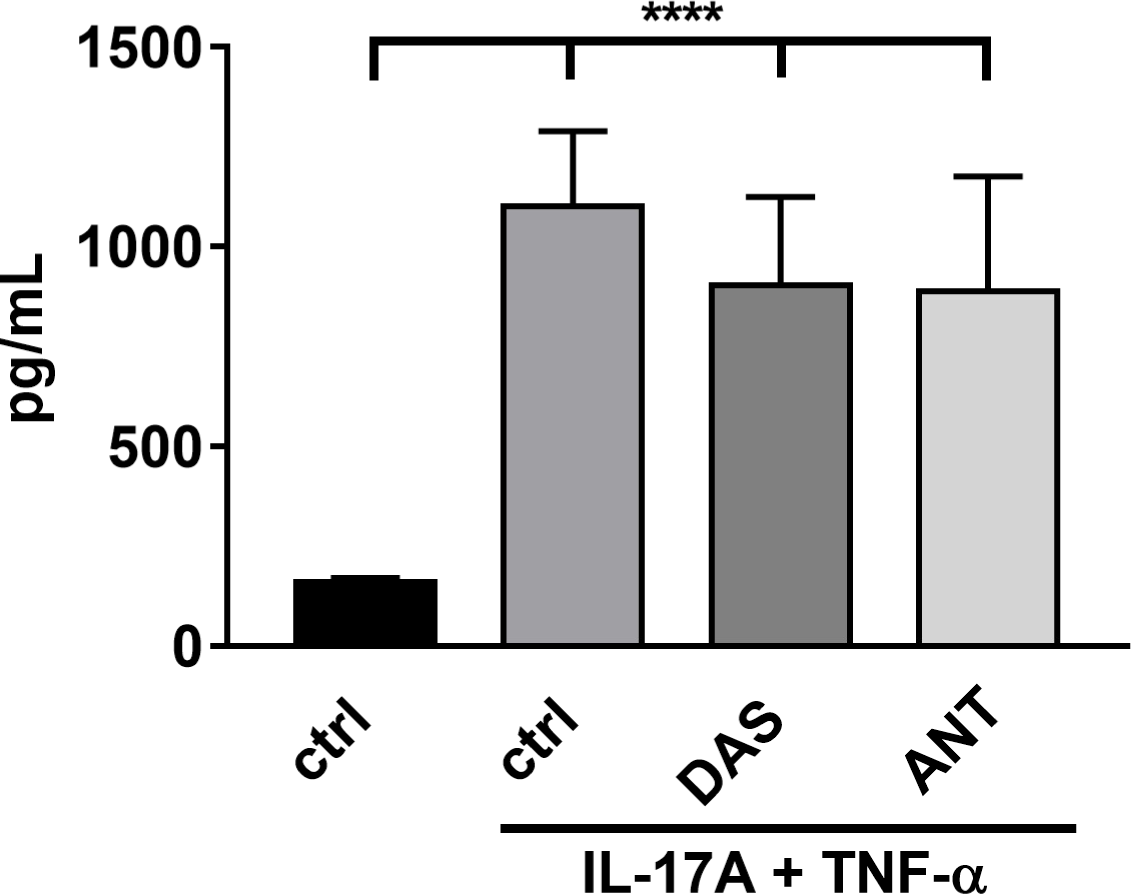
EphA2 inhibition had no effect of secretion of IL-8 induced by IL-17A and TNF-α. IL-8 secretion by oral epithelial cells that were treated with DAS or ANT and then incubated for 20 h with IL-17A (50 ng/mL) and TNF-α (0.5 ng/mL). Results are the mean ± SD of three independent experiments, each performed in duplicate. \*\*\*\**P* < 0.0001.

**Fig. S17.**
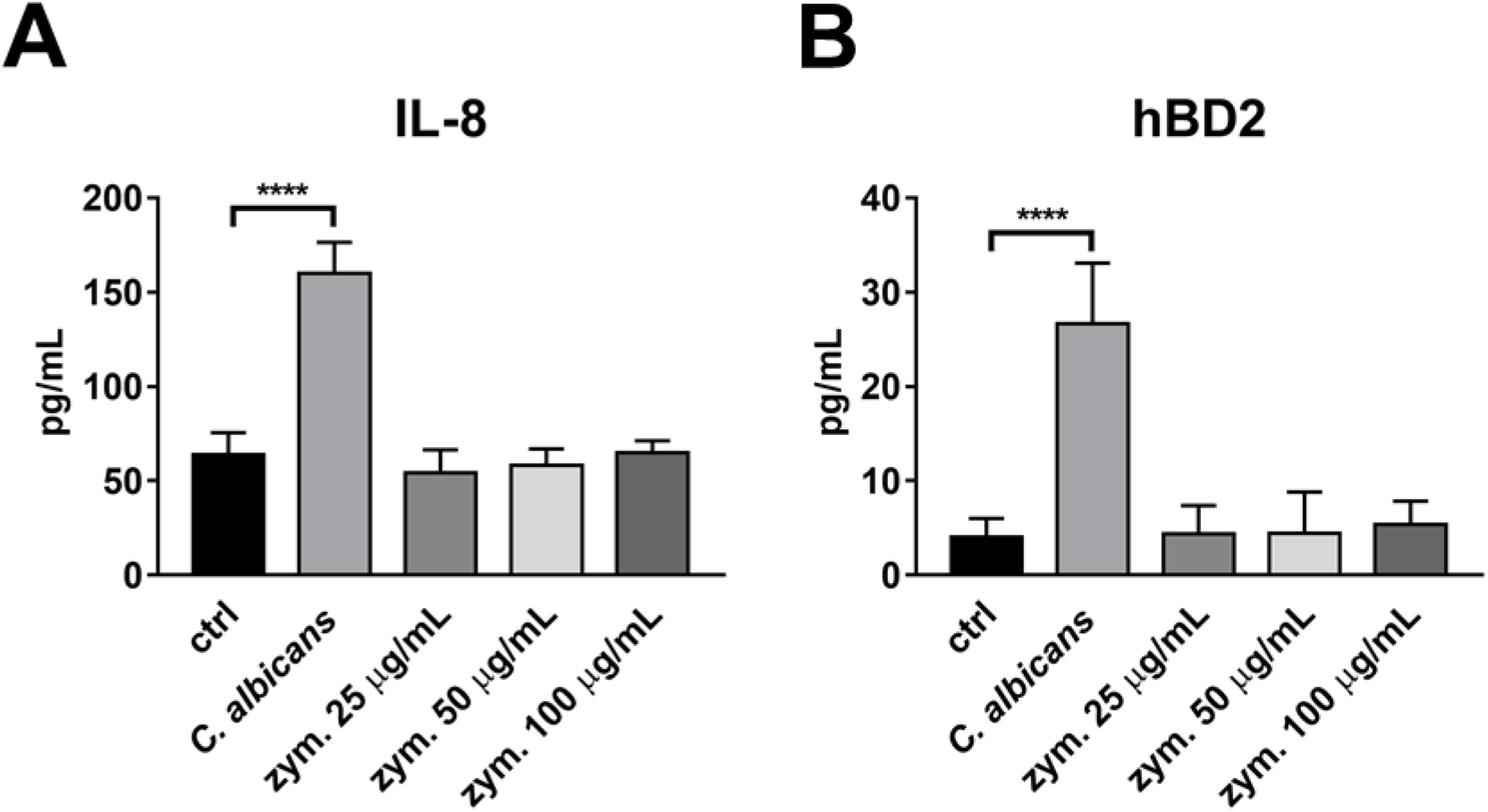
Zymosan does not induces production of IL-8 or hBD2 by epithelial cells. Oral epithelial cells were incubated with *C. albicans* or the indicated concentrations of zymosan, or infected with *C. albicans* for 8 hours and the supernatant were analyzed for IL-8 (**A**) and hBD2 (**B**) content by ELISA. Results are the mean ± SD of three independent experiments, each performed in duplicate. \*\*\*\**P* < 0.0001.

**Fig. S18.**
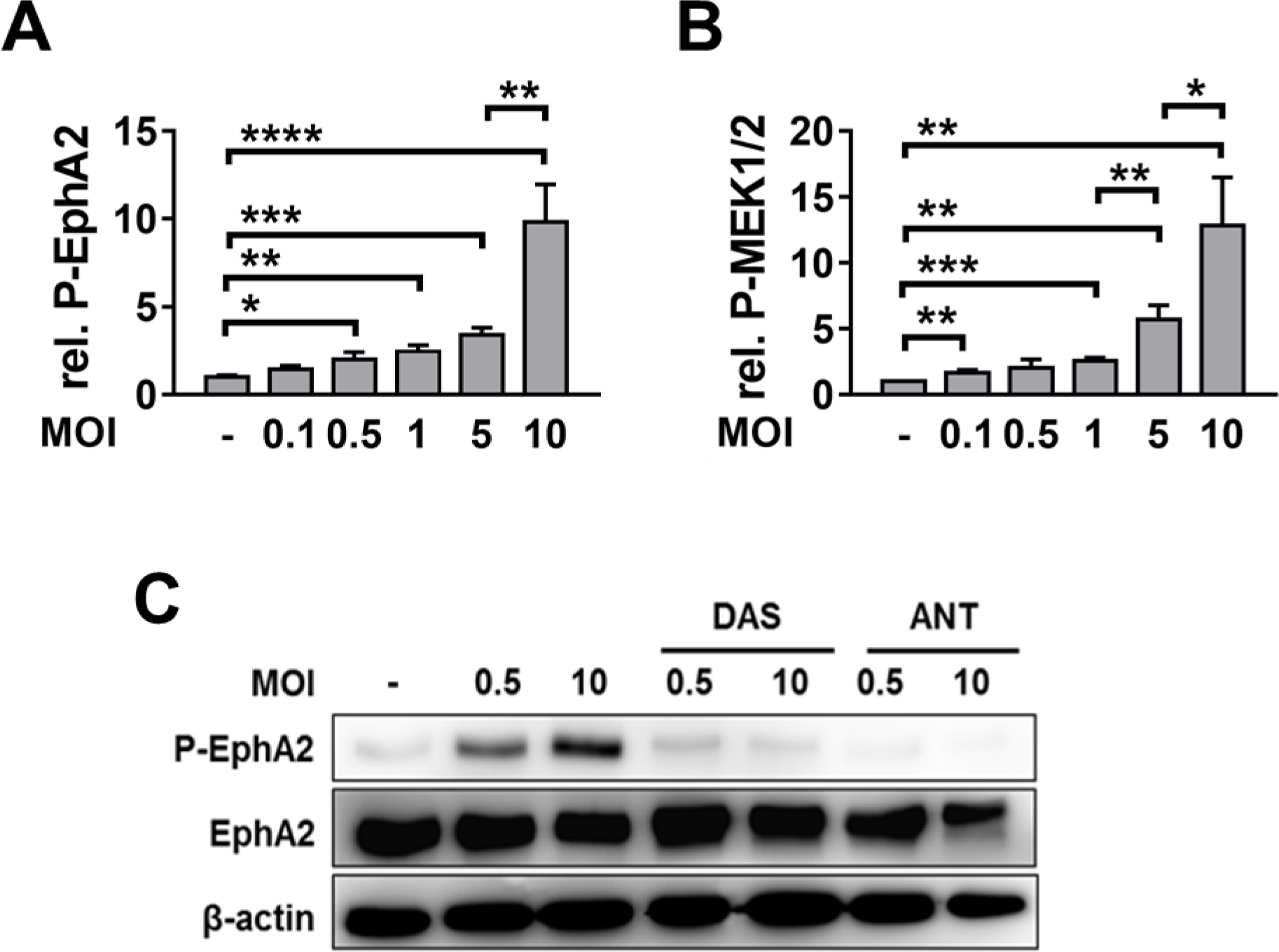
Effects of *C. albicans* inoculum on the phosphorylation of EphA2 and MEK1/2. Densitometric analysis of the phosphorylation of (**A**) EphA2 and (**B**) MEK1/2 in oral epithelial cells that had been incubated for 30 min with *C. albicans* at the indicated multiplicities of infection (MOI). Data are the mean ± SD of 3 independent immunoblots. \**P* < 0.05, \*\**P* < 0.01, \*\*\**P* < 0.001, \*\*\*\**P* < 0.0001. (**C**) DAS and ANT block *C. albicans*-induced phosphorylation of EphA2 at low and high MOIs. Representative immunoblot.

**Fig. S19.**
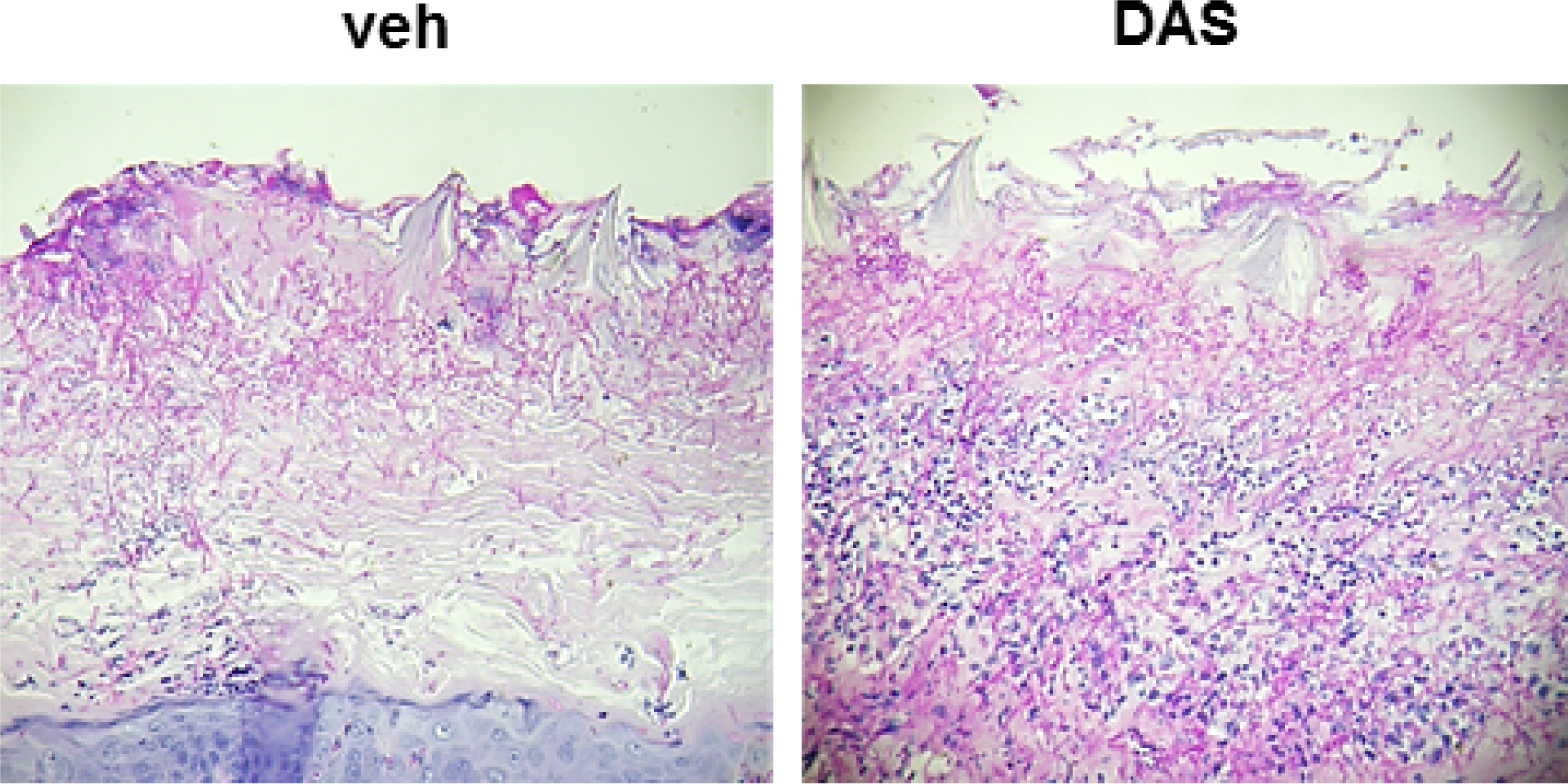
Effects of DAS on the histopathology of the tongues of mice with OPC. Images are of PAS-stained thin sections of the tongues of mice treated with the vehicle control or DAS after 4 d of infection.

**Fig. S20.**
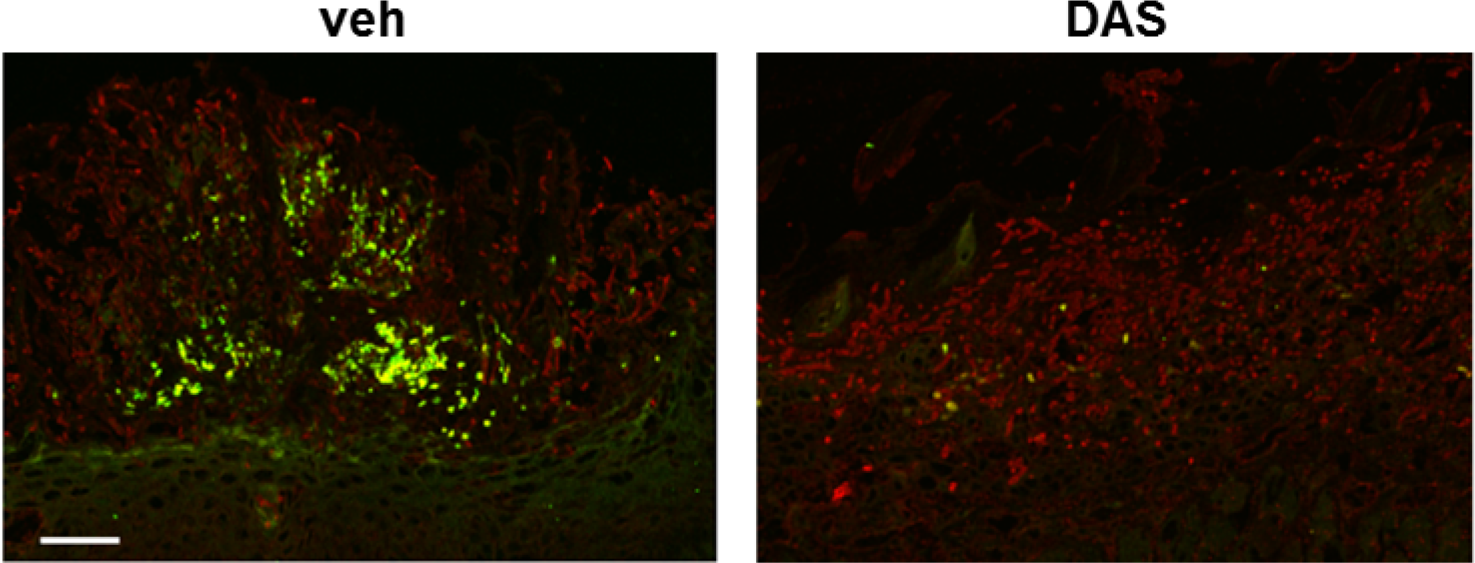
*C. albicans* induces the phosphorylation of Stat3 during OPC. Immunofluorescence images of the tongues of mice treated with the vehicle control or DAS after 4 d of infection. The tongues were stained with antibodies against phospho-Stat3 (green) and *C. albicans* (red). Scale bar 50 μm.

**Fig. S21.**
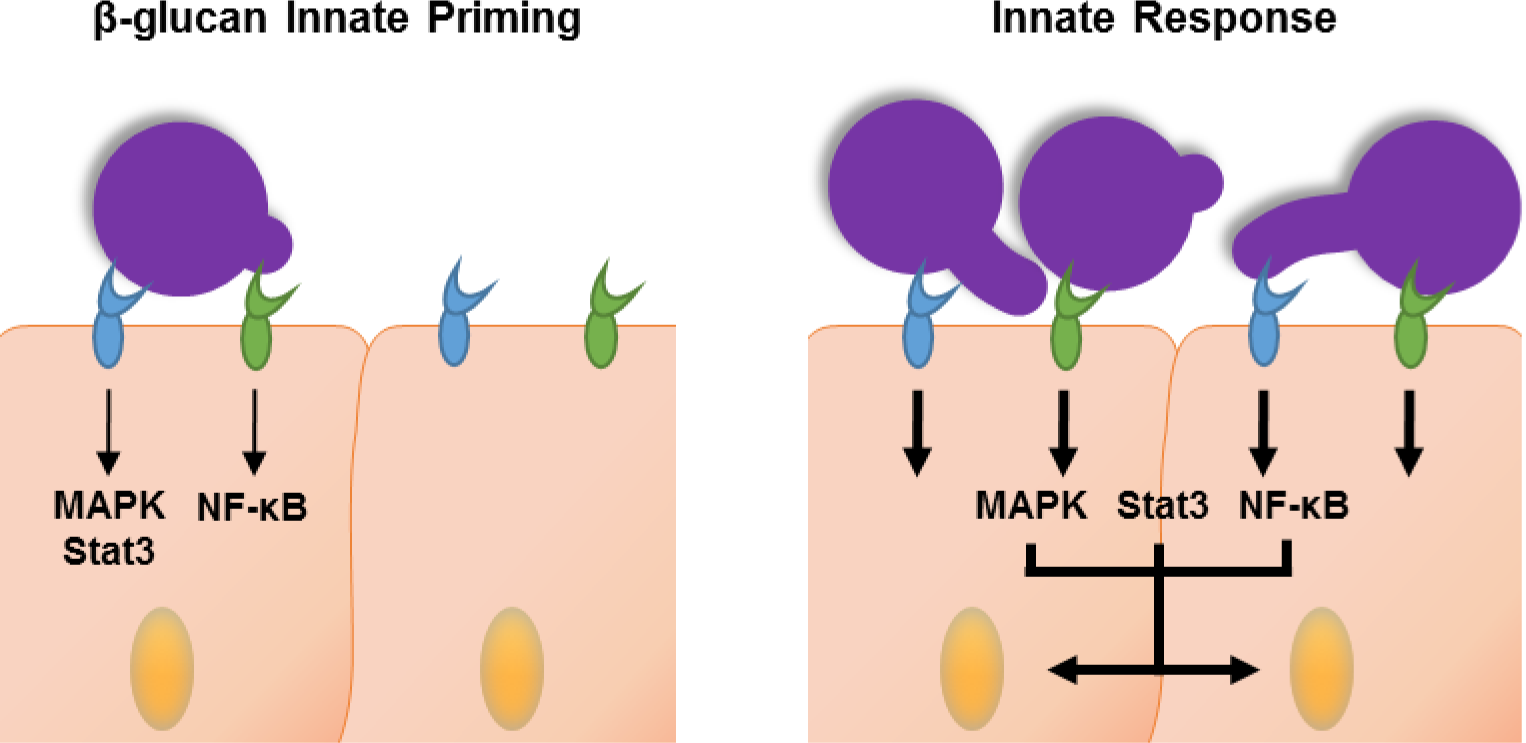
Epha2 on oral epithelial cells binds β-glucan and primes the cells for an inflammatory response. (Left panel) EphA2 (blue) and dectin-1 (green) bind to exposed β-glucan on the fungal surface. Binding to EphA2 activates MAPK and Stat3, whereas binding to dectin-1 activates NF-κB. (Right panel) During fungal proliferation, prolonged activation of EphA2 and possibly dectin-1 induces a pro-inflammatory response in the epithelial cells.

